# Defining effective strategies to integrate multi-sample single-nucleus ATAC-seq datasets via a multimodal-guided approach

**DOI:** 10.1101/2025.04.02.646871

**Authors:** Kathryn Weinand, Erica M. Langan, Michelle Curtis, Soumya Raychaudhuri

## Abstract

**Background:** Chromatin accessibility, measured via single-nucleus Assay for Transposase-Accessible Chromatin with sequencing (snATAC-seq), can reveal the underpinnings of transcriptional regulation across heterogeneous cell states. As the number and scale of snATAC-seq datasets increases, we need robust computational pipelines to integrate samples within a dataset and datasets across studies. These integration pipelines should correct cell-state-obfuscating technical effects while conserving underlying biological cell states, as has been shown for single-cell RNA-seq (scRNA-seq) pipelines. However, scRNA-seq integration methods have performed inconsistently on snATAC-seq datasets, potentially due to sparsity and genomic feature differences.

**Results:** Using single-nucleus multimodal datasets profiling ATAC and RNA simultaneously, we can measure snATAC-seq integration method performance by comparison to independently integrated snRNA-seq gold standard embeddings and annotations. Here, we benchmark 58 pipelines, incorporating 7 integration methods plus 1 embedding correction method with 5 feature sets. Using our command-line tool, we assessed 5 multimodal datasets at 3 different resolutions using 2 novel metrics to determine the best practices for multi-sample snATAC-seq integration. ATAC features outperformed Gene Activity Score (GAS) features, and embedding correction with Harmony was generally useful. SnapATAC2, PeakVI, and ArchR’s iterative Latent Semantic Indexing (LSI) performed well.

**Conclusions:** We recommend SnapATAC2 + Harmony with pre-defined ENCODE candidate *cis*-regulatory element (cCRE) features as a first-pass pipeline given its metric performance, generalizability of features, and method resource-efficiency. This and other high-performing pipelines will guide future comprehensive gene regulation maps.

## Background

Single-nucleus ATAC-seq data (snATAC-seq) defines the open chromatin landscape of individual cells^1^ and has emerged as an important complement to single-cell RNA-seq (scRNA-seq) studies. Open chromatin indicates active regulatory regions such as promoters and enhancers to elucidate the transcriptional regulation of gene expression programs. As scalable snATAC-seq platforms have become affordable and available^2–4^, the number of these datasets has increased dramatically. With this expansion, it is crucial to determine the extent of both shared and distinct cell states, transcription factors (TFs), and ultimately developmental mechanisms^4–6^ and disease therapies^3,7–9^ across datasets and diseases. However, it is challenging to incorporate cells from multiple datasets in different studies^5,6^, encompassing diverse experimental protocols and tissues, as these technical differences often obscure important biological signals. Even in individual large studies^3,5,7^, combining samples obtained across separate batches may result in technical bias. Therefore, effective data integration across snATAC-seq samples and studies is essential.

Technical confounding has also been observed in single-cell and single-nucleus RNA-seq studies^10–14^. By assuming that cellular cell states are shared across subsets of samples, investigators have created highly effective scRNA-seq integration algorithms that remove technical effects while conserving the underlying biology. A recent benchmark^15^ suggested that these scRNA-seq methods effectively balanced conserving biological signals (bio-conservation) and performing batch correction.

However, these same methods have not demonstrated reliable performance for snATAC-seq data^15^, perhaps due to differences in genomic features. While scRNA-seq analysis initially uses a pre-defined set of ∼20,000 protein-coding genes, snATAC-seq analysis does not have a universal feature set. Peaks of localized signals called within a dataset are the most common feature type for snATAC-seq data. However, snATAC-seq data typically has fewer reads mapping to a larger feature set (∼100,000 peaks)^16^ than scRNA-seq data. Therefore, single cell integration methods, largely derived for scRNA-seq data, struggle with the increased sparsity when applied to snATAC-seq data.

To assess bio-conservation and batch correction for a snATAC-seq integration pipeline, a benchmarking study requires gold standard datasets with clear ground truths. Previous snATAC-seq integration benchmark studies have used datasets with cell types defined using the tested modality^15^, with broad cell type annotations that are relatively easy to distinguish^15^, with confounded cell types and batches^17^, or avoided batch effects altogether^18^. However, we can strategically utilize multimodal datasets that simultaneously capture gene expression and chromatin accessibility measurements for the same cells. In these instances, we can define a gold standard cell type and embedding by integrating snRNA-seq profiles with state-of-the-art methods shown to be effective in both bio-conservation and batch correction^15,19^. This embedding can be used as an independent high-resolution standard to benchmark snATAC-seq integration pipelines. To assess the quality of snATAC-seq integration, we use metrics assessing integration at the most local level by comparing to the independent snRNA-seq integrated embeddings for the same cells.

Here, we benchmarked 58 snATAC-seq data integration pipelines across 7 methods, 5 feature sets, and 1 embedding correction^20^ to define best practices for the genomics community (**Fig. 1a-b**; **Additional file 1: Table S1**). We assessed each pipeline using 2 novel local metrics (**Fig. 1c-f**) to quantify efficacy in integration while preserving subtle functional differences between cells. We also used 2 established metrics for comparison. Using our command-line tool, we applied these strategies across 5 diverse blood and tissue datasets^21,22^ at increasing resolutions that represent increasing integration difficulty (**Fig. 1g-h**). Overall, SnapATAC2^23^ + Harmony^20^ using a pre-defined set of ENCODE candidate *cis*-regulatory element^19^ (cCRE) features had the best performance. ArchR^24^ and PeakVI^25^ were also among the better-performing methods, though the latter was more resource-intensive. Furthermore, we found that ATAC features (peaks, cCREs, tiles) consistently performed better than Gene Activity Score (GAS) features.

**Fig. 1.**
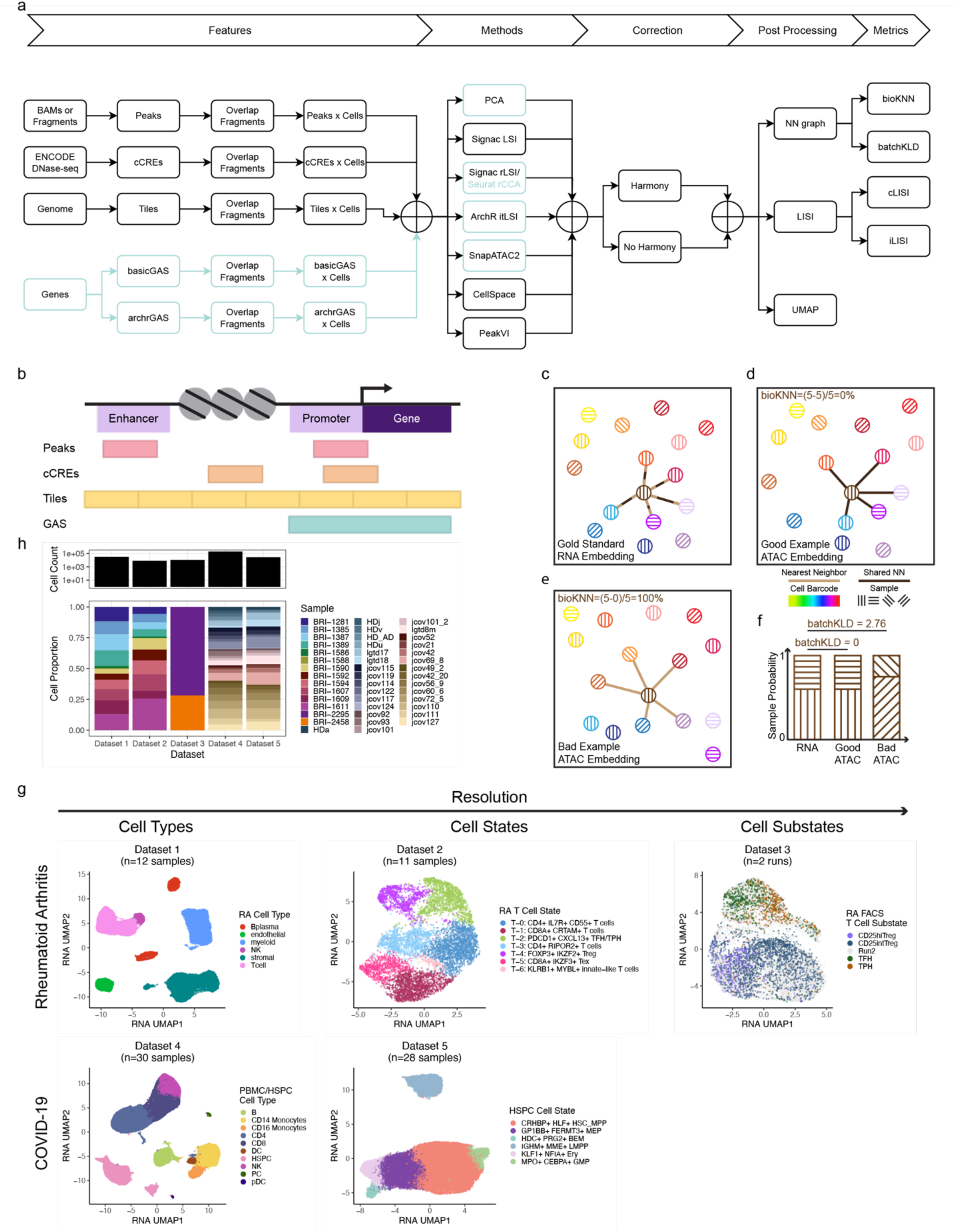
Benchmark design. **a.** Overview of snATAC-seq integration benchmark. GAS features only used methods highlighted in blue and used Seurat rCCA instead of Signac rLSI (**Methods**). **b.** For an example locus, a visual comparison of the four feature types tested: peaks, cCREs, tiles, GASs. **c.-e.** bioKNN metric. Cell identity is denoted by color. **c.** For the brown cell, the nearest neighbors (NN) defined by snRNA-seq. NN edges are striped to correspond with both **d.** and **e.**. **d.** In a good example, the brown cell’s NN defined by snATAC-seq; since all top 5 NN are shared in both modalities, denoted by dark brown NN edges, the bioKNN is (5-5)/5=0%, the best possible value. **e.** In a bad example, the brown cell’s NN defined by snATAC-seq; since no top 5 NN are shared in both modalities, denoted by tan NN edges, the bioKNN is (5-0)/5=100%, the worst possible value. **f.** Batch Kullback-Leibler Divergence (batchKLD) metric. Sample identity is denoted by pattern. The batchKLD between top NN from **c.** and **d.** was 0, the best possible value. The batchKLD between top NN from **c.** and **e.** was 2.76, this example’s worst possible value (**Methods**). **g.** We tested five multimodal datasets from two studies^21,22^: (1) RA inflammatory tissue broad cell types, (2) RA inflammatory tissue T cell states, (3) RA PBMC T cell substates, (4) COVID-19 PBMC broad cell types, and (5) COVID-19 PBMC HSPC states.

## Results

### Benchmark Design

We benchmarked pipelines consisting of three parts: one of five feature sets, one of seven integration methods, and the presence or absence of a correction method (**Additional file 1: Table S1**); this was followed by post-processing and quantification of performance metrics (**Fig. 1a**). The seven integration methods used the feature matrices to create an initial embedding while the correction method, Harmony^20^, adjusted those embeddings to account for batch effects.

Dataset 3 cell phenotypes were determined using FACS (**Methods**). Otherwise, the cell phenotypes were determined by multimodal snRNA-seq. UMAPs were defined using snRNA-seq. **h.** The total (log10) cell counts across all 5 Benchmarking Datasets (**top**) as well as the proportion of cells per sample (**bottom**). The samples in Datasets 4/5 are grouped by healthy (blue), non-COVID-19 (slate), mild/mild_late COVID-19 (pinkish gray), early COVID-19 (pink), and late COVID-19 (yellow).

Of the 5 different feature sets, 3 were ATAC-specific features and 2 were gene-like GAS features (**Fig. 1b**; **Table 1**). Peaks called using each dataset’s BAM or fragment files were the most dataset-specific feature set; these features described the intrinsic signal within a dataset but transfer poorly to other datasets. In contrast, tiles, non-overlapping fixed-length sliding windows tiled across the entire genome, were highly transferrable. Tiles, however, were memory-intensive, with a set size an order of magnitude larger than called peaks. Furthermore, arbitrary tile boundaries may disrupt meaningful biological signals. cCREs were regulatory regions called from thousands of ENCODE bulk DNase-seq experiments spanning diverse cell types^19^; they had the generalizability of tiles, but the biological value of peaks, though they may miss some dataset-specific biology. We fixed each of these three snATAC-seq features to be 500 bp to avoid length as a confounding factor (**Methods**). For the GAS features, we aggregated signal in and around genes in two different ways. The “basicGAS” score simply counted fragments that overlapped the 2 kb promoter and gene body. Increasing complexity, “archrGAS” score used ArchR’s^24^ default exponential decay model, which weights tiles overlapping gene windows, with high weights in the 5 kb promoter and gene body, but decreased weights further away; it was also linearly scaled for gene length. As with genes, GASs were transferable across datasets. We note that since linking of ATAC reads to target genes remains challenging, GAS scores may not accurately reflect the biological signal of a gene. Indeed, within broad cell types using multimodal snATAC-/snRNA-seq data, we previously^26^ found that there was minimal correlation between real gene expression and five different GAS methods, including the two used here.

**Table 1.** Feature summary.

| Feature | Peak | cCRE | Tile | GAS |
| --- | --- | --- | --- | --- |
| Dataset-specific features | Yes | No | No | No |
| Localized signal | Yes | Maybe | No | No |
| Biologically meaningful | Yes | Yes | No | Yes |
| Transferrable | No | Yes | Yes | Yes |
| Standard size | Enforced | Enforced | Yes | No |
| Feature counts | ~100K | ~600K | ~6M | ~20K |
| Overlapping features | Yes | Yes | No | Yes |
| Tn5 bias correction | Yes | No | No | No |

cCRE: candidate cis-regulatory element; GAS: gene activity score.

We assessed seven integration methods for initial embedding generation: (1) Principal Component Analysis (PCA), (2) Signac^27^ using latent semantic indexing (LSI), (3) Signac using reciprocal LSI (rLSI), (4) ArchR^24^ using iterative LSI (itLSI), (5) SnapATAC2^23^ (SA2) using matrix-free spectral embedding, (6) CellSpace^28^ (CS) using neural embedding, and (7) PeakVI^25^ (pVI) using a variational autoencoder (**Table 2**). These methods reflected linear (1-4) and non-linear (5-7) options. Only rLSI/rCCA and PeakVI specified a batch variable for explicit batch correction. Since CellSpace, PeakVI, Signac LSI, and Signac rLSI were implemented for use with ATAC features, we did not test GAS features with them. However, to test GAS features in a framework similar to Signac, we used Seurat^29^, a method originally designed to analyze scRNA-seq data, with GAS features inputs. We specifically tested Seurat’s batch-aware reciprocal canonical correlation analysis (rCCA) method, which is related to the anchor integration approach used by rLSI.

**Table 2.**
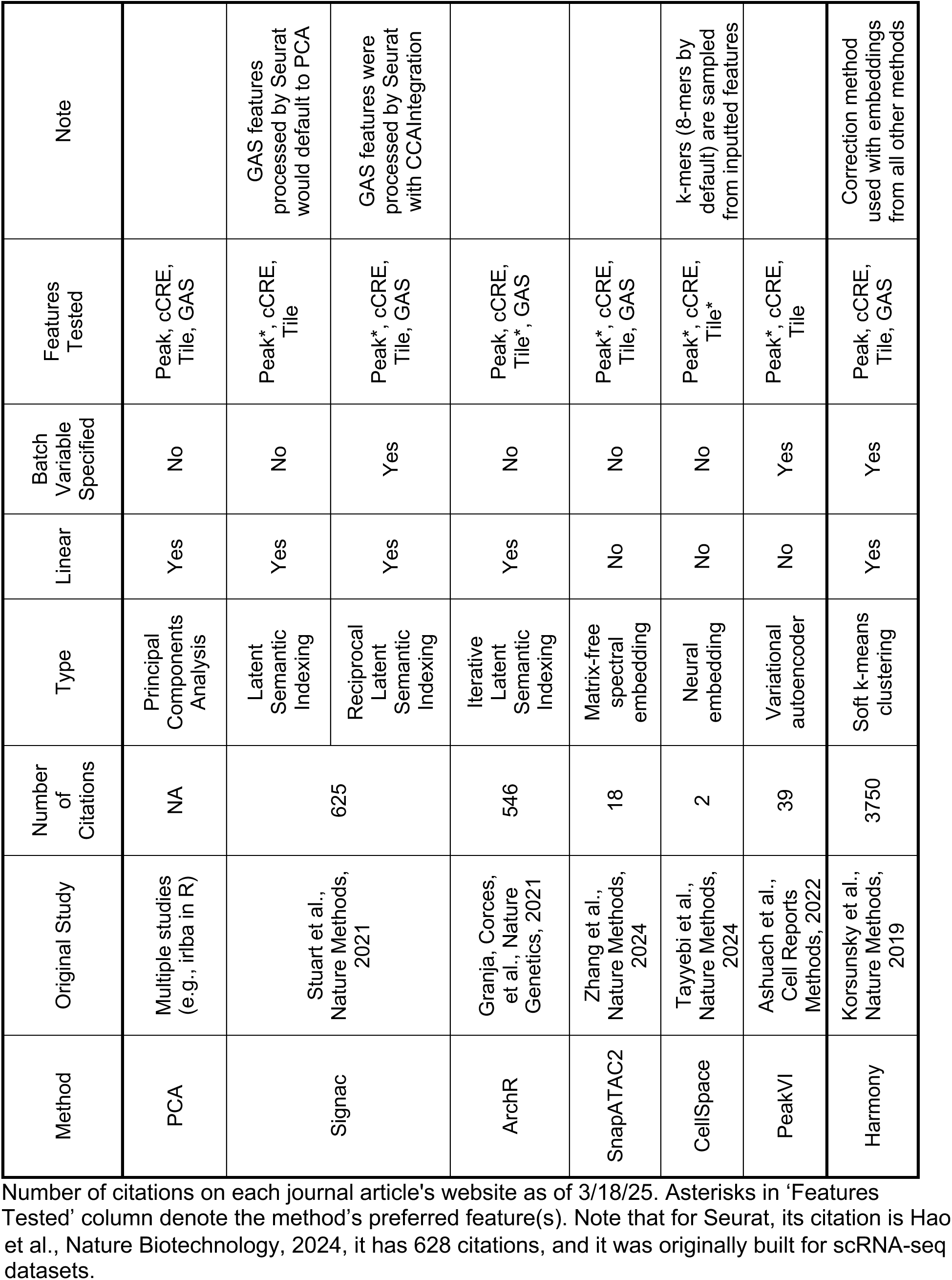
Methods summary.

To assess the value of additional batch correction, we tested the inclusion of a high-performing^15^ embedding correction method, Harmony^20^ (**Table 2**). While Harmony was initially developed for scRNA-seq data, it is often used in conjunction with the assessed snATAC-seq methods^7,21,22^. Harmony corrected each previously-defined embedding to integrate samples within a dataset. We denoted each embedding without Harmony as “Method” and its Harmony-corrected embedding as “Method + Harmony”.

We assessed each snATAC-seq embedding by comparing it to a gold-standard embedding using two per-cell nearest-neighbor (NN) metrics we developed: bio-conservation KNN (bioKNN) and batch Kullback-Leibler divergence (batchKLD). We chose local-based metrics rather than cluster-based metrics since the latter can obscure inaccuracies by aggregation. Our metrics relied on multimodal data, which provided two readouts, ATAC and RNA, of the same underlying biological identity. Assuming modality-specific batch differences were adequately addressed, each cell should be surrounded by similar cells in both modalities. These shared cells can be identified by cell barcodes present in both modalities, thus avoiding the need for cell type annotations. Using well-established scRNA-seq integration methods, we first created a gold standard embedding from the multimodal RNA, upon which we defined the NN for each cell (**Fig. 1c**). Then, for each snATAC-seq integration strategy, we used its embedding to define per-cell NN (**Fig. 1d-e**). Finally, for each cell, we counted how many of its ATAC top K NN overlapped with its RNA top K NN and quantified bioKNN as the percentage of K cells that were not shared across modalities (**Methods**). A lower bioKNN value reflected a more aligned integration where the underlying biology was shared across modalities.

Furthermore, since all the shared cells could be from the same sample and hence poorly integrated, we also assessed batch correction using batchKLD (**Fig. 1f**) between the per-cell sample distributions of the top K NN for the gold standard RNA embedding (**Fig. 1c**) and the top K NN for each ATAC embedding (**Fig. 1d-e**). A lower batchKLD value was favorable, meaning the sample distributions between the two modalities were more similar; the upper bound was blunted by a dataset-driven pseudocount (**Methods**).

For comparison, we also quantified commonly used local benchmarking metrics^15,18,23,29^, introduced in Korsunsky et al.^20^, defined by Local Inverse Simpson’s Index (LISI) scores: cell type LISI (cLISI) for bio-conservation and integration LISI (iLISI) for batch correction. A low cLISI score was preferable, signifying that cell types were not mixing and biological variation was preserved. A high iLISI score indicated more sample mixing and suggested an effective correction of sample-specific technical variation. While useful, these metrics had some pitfalls addressed by our NN metrics. Unlike the bioKNN metric, cLISI required cell type annotations, which are often challenging to define in an orthogonal manner. iLISI assumed uniform sample mixing for each cell type was desired, which may not be true in all biological contexts: for example, a cell type might be missing in healthy controls but present in disease samples. batchKLD used the integrated snRNA-seq sample distribution to account for such instances. We calculated all four metrics for each cell.

## Datasets tested

We tested each of the 58 strategies on 5 different 10x multiome datasets (**Fig. 1g-h**; **Table 3**), for a total of 290 snATAC-seq embeddings. Since the resolution changed by dataset, we denoted cell types, states, and substates collectively as “cell phenotypes.” As the resolution increased and cell phenotypes became more similar to each other, it should become harder for these pipelines to distinguish between them. Using each dataset’s quality-controlled cell set (**Additional file 2: Fig. S1-5a-b**), we reprocessed the snRNA-seq data from feature selection to UMAP for uniformity (**Additional file 2: Fig. S1-5c-d**; **Methods**). We reassigned gold standard biological cell phenotypes with a PCA + Harmony mRNA clustering resolution that most closely resembled the original author-defined annotations (**Methods**; **Additional file 2: Fig. S1-5e-i**).

**Table 3.** Dataset summary.

| Benchmarking Dataset | Original Study | Source | Cell Phenotype | Cell Count | Sample Count |
| --- | --- | --- | --- | --- | --- |
| 1 | Weinand et al., Nat Commun, 2024 | RA & OA inflammatory tissue | Broad cell types | 31,547 | 12 |
| 2 | Weinand et al., Nat Commun, 2024 | RA & OA inflammatory tissue | T cell states | 8,069 | 11 |
| 3 | Weinand et al., Nat Commun, 2024 | RA PBMC | T cell substates | 10,669 | 2 runs of 4 pooled samples each |
| 4 | Cheong et al., Cell, 2023 | COVID-19 & healthy PBMC | Broad cell types | 197,360 | 30 |
| 5 | Cheong et al., Cell, 2023 | COVID-19 & healthy PBMC | HSPC states | 27,979 | 28 |

RA: rheumatoid arthritis; OA: osteoarthritis; PBMC: peripheral blood mononuclear cell; COVID-19: coronavirus disease 2019; HSPC: hematopoietic stem and progenitor cell

Within that re-processing, we generated NN graphs for use in our NN metrics for primary analysis. To assess sensitivity of our NN metrics across snRNA-seq integration pipelines, we also generated a Seurat rCCA RNA embedding for a secondary analysis (**Methods**). We excluded samples with fewer than 100 cells since they did not perform well with Seurat rCCA default parameter settings. For primary analysis, we generated NN graphs with 200 cells, though we also used 50 and 100 neighbors to confirm trends.

The first three datasets originated from our previous study^21^ investigating rheumatoid arthritis synovial tissue and spanned the gamut from broad cell types (“Dataset 1”) to T cell states (“Dataset 2”) to T cell substates (“Dataset 3”) (**Fig. 1g**). Dataset 1 had 12 samples for a total of 31,547 cells while Dataset 2 totaled 8,069 cells after dropping a sample for low cell count (**Fig. 1h**). One sample in each came from an osteoarthritis patient. Dataset 3 was of particular interest since its cell phenotypes were determined via Fluorescence-Activated Cell Sorting (FACS) surface markers: CD4^+^CD127^-^CD25^hi^ regulatory T cells (Treg), CD4^+^CD127^-^ CD25^int^ Treg, CD4^+^CD25^-^PD1^+^CXCR5^+^ T Follicular Helper cells (TFH), and CD4^+^CD25^-^ PD1^+^CXCR5^-^ T Peripheral Helper cells (TPH). While mRNA clusters aligned well to the protein hashtags and differential genes were found between the two Treg substates and between the TFH/TPH substates in the original study, we were unable to find many differential promoter peaks^21^. Thus, it should be the most challenging to phenotypically characterize in snATAC-seq embeddings. However, it did have the fewest number of batches with 2 runs of 4 pooled samples each, for a total of 10,669 cells (**Fig. 1h**; **Methods**).

The last two datasets came from Cheong et al., 2023^22^, where the authors applied a newly developed method for peripheral blood mononuclear cell analysis with progenitor input enrichment (PBMC-PIE). They used this method to profile both mature immune cell types (“Dataset 4”) and hematopoietic stem and progenitor cells (HSPCs; “Dataset 5”) from 30 PBMC samples across 5 patient stratifications: healthy, non-COVID-19 critical illness, mild COVID-19, early COVID-19, and late COVID-19 (**Methods**; **Fig. 1g-h**). Of note, Dataset 4 stress-tested the time and memory limits for our pipelines as it had almost 200K cells. It was generally more challenging to determine cell phenotypes in Dataset 5 for its 27,979 progenitor cells instead of fully differentiated cell states.

### Benchmarking 58 different pipelines

Using our command-line tool, we ran each of the 58 pipelines on the five datasets and quantified metrics for bio-conservation and batch correction (**Additional file 2: Fig. S6**).

To illustrate results, we focused on Dataset 5 (**Fig. 2a**). We observed specific pipeline choices affected bio-conservation and batch correction. Generally, the best-performing pipelines for this dataset used SnapATAC2 or itLSI. GAS features predominately resulted in poor bio-conservation while the inclusion of Harmony usually improved batch integration. For example, SnapATAC2 + Harmony with peaks generally performed very well (**Fig. 2b**), with distinct cell states and diverse samples that corresponded well with the snRNA-seq data (**Fig. 1g**; **Additional file 2: Fig. S5c**). However, replacing peak features with archrGAS features resulted in a similar mean batchKLD metric, but a worse mean bioKNN metric (**Fig. 2a**); this was reflected in the UMAP where cell states were mixed together despite comparable sample diversity (**Fig. 2c**). Conversely, using SnapATAC2 with peaks but removing the Harmony correction step resulted in a worse mean batchKLD metric (**Fig. 2a**); in the UMAP, there were groups of cells originating from a single sample (**Fig. 2d**) that had correspondingly high batchKLD metrics (**Fig. 2e**) since the snRNA-seq data did not contain these singular sample regions (**Additional file 2: Fig. S5c**). Including Harmony largely did not change the bio-conservation metric, with the best bioKNN values in the smaller cell states, BEM and GMP (**Fig. 2e**).

**Fig. 2.**
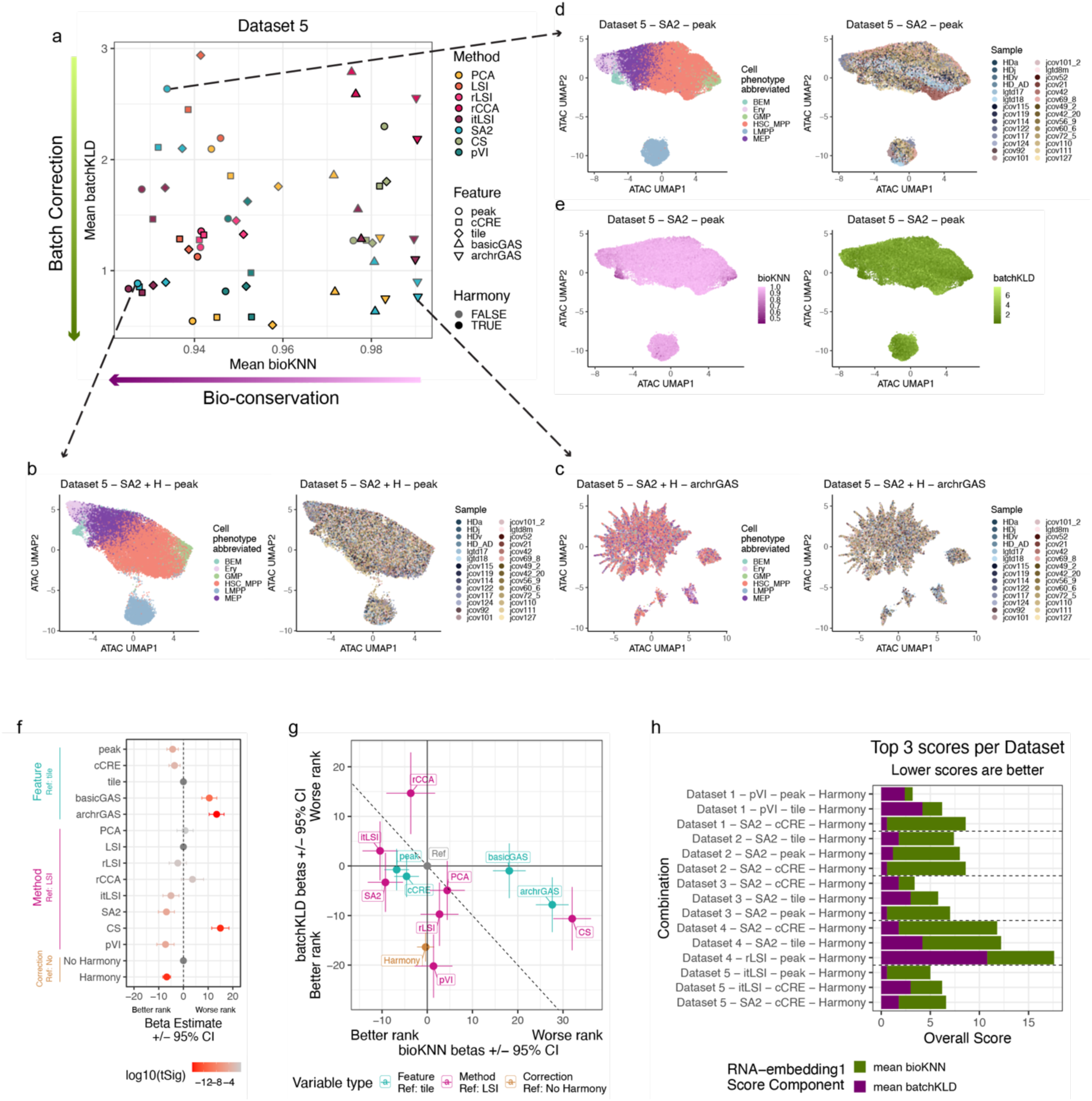
Pipeline rankings using NN metrics with RNA-embedding1. **a.** Mean bio-conservation (**x-axis**; bioKNN) and batch correction (**y-axis**; batchKLD) metrics averaged across Dataset 5 cells. **b.**-**d.** snATAC-seq UMAPs colored by cell phenotype (**left**) and sample (**right**) for Dataset 5 cells using **b.** SnapATAC2 + Harmony with peak features, **c.** SnapATAC2 + Harmony with archrGAS features, **d.** SnapATAC2 with peak features. **e.** snATAC-seq UMAPs colored by bioKNN (**left**) and batchKLD (**right**) for Dataset 5 cells using SnapATAC2 with peak features. In both cases, lower values (darker colors) are desired. **f.** Combined linear model relating overall score (60% ranked mean bioKNN + 40% ranked mean batchKLD) to dataset, feature, method, and correction combinations (**Methods**). **g.** Separated linear models as in **f.**, but ranked mean bioKNN on x-axis and ranked mean batchKLD on y-axis (**Methods**). **h.** Top 3 overall scores per dataset, colored by ranked mean bioKNN and ranked mean batchKLD components.

### General trends across five datasets

To understand the overall trends, we examined the mean bio-conservation and batch correction metrics across all 58 pipelines and 5 datasets. For each dataset, we ranked performance (1 best; 58 worst) for each of the pipelines and examined a composite rank of bio-conservation and batch correction, weighted by 60% and 40%, respectively^15^ (**Methods**); we also ranked the individual metrics. We assessed overall performance using a multivariate linear model, where we determined how features, methods, and embedding correction affected the ranked mean scores across the five datasets^15^ (**Fig. 2f**). We denoted our primary analysis focusing on NN metrics utilizing 200 NN within an embedding generated from snRNA-seq datasets using PCA with Harmony correction, as RNA-embedding1. We also did secondary analyses with NN metrics using Seurat gene embeddings and 200 NN (‘RNA-embedding2’) (**Additional file 2: Fig. S7a**) and LISI metrics (**Additional file 2: Fig. S7b**).

First, we found that feature set (p<0.00227) impacted performance in the following order: archrGAS (worst; beta= +13.45 rank), basicGAS, tile (reference rank=0), cCRE, peak (best; beta= -4.36). Second, across datasets and pipelines, the choice of method nominally affected rank performance (p<0.12). We observed that rLSI, itLSI, SnapATAC2, and PeakVI improved rank performance when compared to the reference LSI (beta= -2.27, -5.05, -6.89, -7.26, respectively). Globally, SnapATAC2 was hurt by the GAS features that were not included in the PeakVI feature sets tested. In contrast, PCA had little impact on the rank (beta= +0.67) while rCCA had worsened ranks (beta= +3.67). CellSpace had the worst rank performance and the biggest effect size across methods (beta= +14.97). Third, the decision to include Harmony as an additional batch correction step generally improved the overall rank performance (p=3.36e-15; beta= -6.77). We observed similar patterns with NN metrics with RNA-embedding2 and LISI metrics (**Additional file 2: Fig. S7a-b**). A notable exception was for Signac rLSI, where it only became significantly different from the LSI reference in the NN metrics using RNA-embedding2 defined from Seurat rCCA (p=0.00025) compared to the RNA-embedding1 (p=0.20) and LISI (p=0.16) metrics, suggesting that the choice of RNA embedding may have a subtle impact. Seurat rCCA results also changed minimally across metric types, but their ranks were already penalized by the worst-preforming GAS features.

To understand the effect of pipeline choices on bio-conservation and batch integration separately, we applied the same approach, but testing a linear model for each metric (**Methods**; **Fig. 2g**; **Additional file 2: Fig. S7c-d**). We observed that the choice of feature set had the greatest effect on bio-conservation metrics (p<0.0011). SnapATAC2 was the only method to achieve better rankings for both bio-conservation and batch correction for both NN metrics. Unsurprisingly, Harmony primarily improved batch correction metrics (p=1.09e-23), as described in the Dataset 5 analysis above (**Fig. 2b,d**).

### ATAC-specific features surpassed Gene Activity Scores

One of the most striking results of this study was that ATAC-specific features, rather than GAS scores, generally improved performance across pipelines and metrics (**Fig. 2f**; **Additional file 2: Fig. S7a-b**). None of the top 3 combinations per dataset included the GAS features (**Fig. 2h**; **Additional file 2: Fig. S7e-f**). Indeed, using the RNA-embedding1 NN metrics, the best pipeline containing GAS features was Dataset 2 using itLSI + Harmony with basicGAS features (score=25.2); it did not surpass the first quartile of scores for that dataset (score=20.45; **Additional file 1: Table S2**). Consistent with this observation, the corresponding UMAPs illustrated inappropriate mixing between the two primarily CD8+ T cell populations (T-1 and T-5) and between the TFH/TPH and Treg populations (T-2 and T-4) while showing pockets of cells segregated from two samples (**Fig. 3a, left**).

**Fig. 3.**
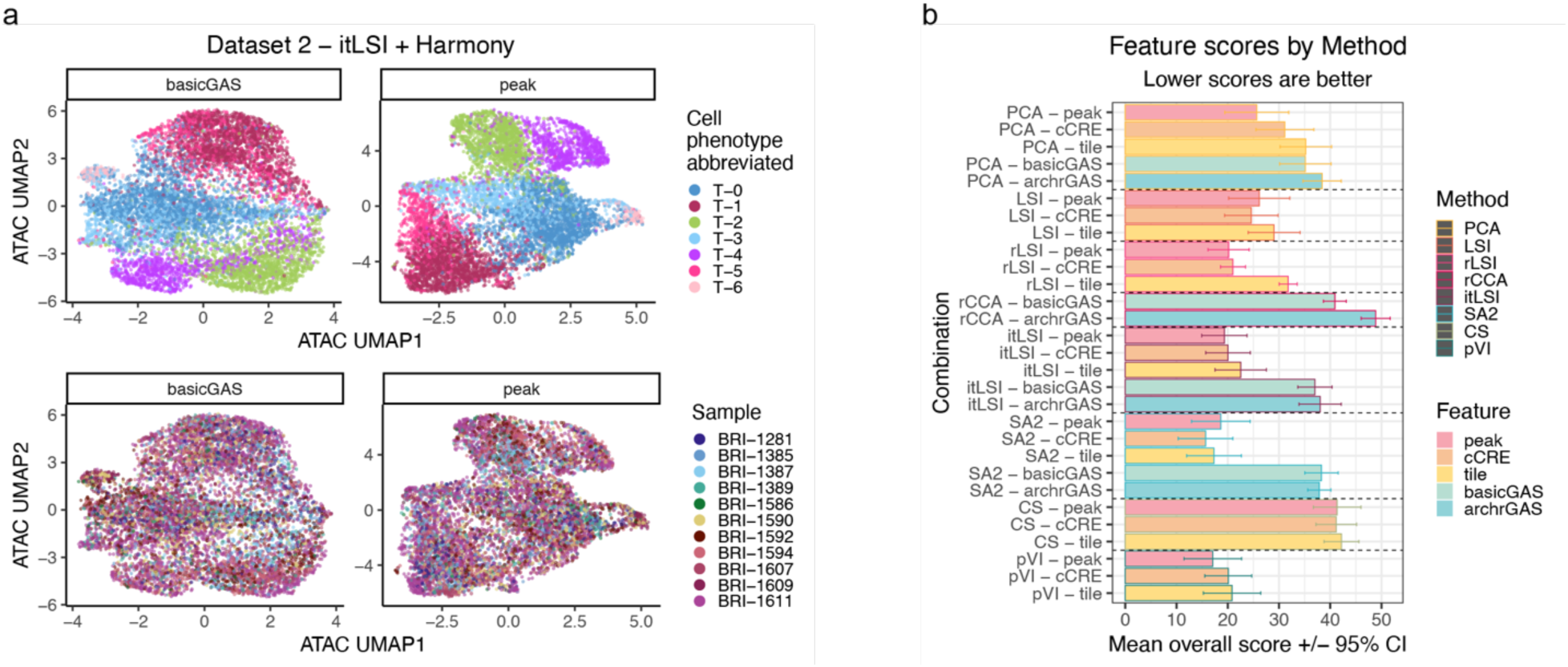
ATAC features (peaks, cCRE, tiles) outperformed GAS features (basicGAS, archrGAS). **a.** Best performing GAS feature pipeline (**left**): Dataset 2 using basicGAS features and the itLSI method with Harmony correction. Substituting peak features within that pipeline (**right**). Corresponding UMAPs colored by cell phenotype (**top**) and sample (**bottom**). **b.** The mean overall scores (60% bio-conservation bioKNN rank + 40% batch correction batchKLD rank) across datasets and correction choice for each method and feature combination. Lower scores are better. Error bars are the 95% confidence interval (CI).

Amongst ATAC features, we generally observed that peaks and cCREs yielded the best overall performance, followed closely tiles (**Fig. 2f**). This pattern was generally consistent across methods (**Fig. 3b**; **Additional file 2: Fig. S8a-b**). For the above Dataset 2 itLSI + Harmony pipeline, simply switching the basicGAS features for peak features both improved the score (12.4) and resulted in more visually distinct T cell states (**Fig. 3a, right**). We note that peaks are the most commonly used snATAC-seq feature, and most of these methods were likely developed assuming peak features. That being said, the majority of dataset-specific peaks (mean=70%) overlapped at least one cCRE (**Additional file 2: Fig. S8c**), perhaps accounting for the generally small differences in metric rankings between them. While tiles overlapped with all other features since tiles accounted for the whole genome, they were not centered at the summit of accessibility unlike the peaks (**Methods**).

### SnapATAC2 performed best using NN metrics

Using the RNA-embedding1 NN metrics, SnapATAC2 had the best-ranked combination for 3/5 datasets, and was among the top 3 for the other 2 datasets (**Fig. 2h**). Its success was most apparent in the bio-conservation metrics (**Fig. 2g**). While all ATAC features performed well for SnapATAC2, it did best with cCREs (**Fig. 3b**, **4**; **Additional file 1: Table S2**). SnapATAC2 could handle GAS features with limited modification (**Methods**), but as with all other methods, the GAS features hurt performance (**Additional file 2: Fig. S6a**; **Additional file 1: Table S2**). SnapATAC2 + Harmony with ATAC features generally performed well when using the RNA-embedding2 NN metrics as well (**Additional file 2: Fig. S6b**).

### PeakVI performed well in the rheumatoid arthritis datasets, but biased by fragment counts

In every dataset, we observed that PeakVI + Harmony used with peaks outperformed all other PeakVI pipelines via the RNA-embedding1 NN metrics (**Additional file 1: Table S2**). In fact, this combination was the best-performing pipeline for Dataset 1 (**Fig. 2h**). In the corresponding UMAP, NK cells separated from T cells and mural cells segregated from stromal cells in the lower tail (**Fig. 5a**). It also performed well for Datasets 2 and 3, primarily owing to effective batch correction (**Fig. 2g**; **Additional file 1: Table S2**). However, PeakVI’s performance decreased in the COVID-19 datasets, due to worse bio-conservation rankings (**Additional file 1: Table S2**). PeakVI struggled to distinguish pDCs in Dataset 4 and BEM in Dataset 5 (**Fig. 5a**). It also displayed some mixed HSPC states for the late COVID-19 samples in Dataset 4 (**Fig. 5b**).

**Fig. 4.**
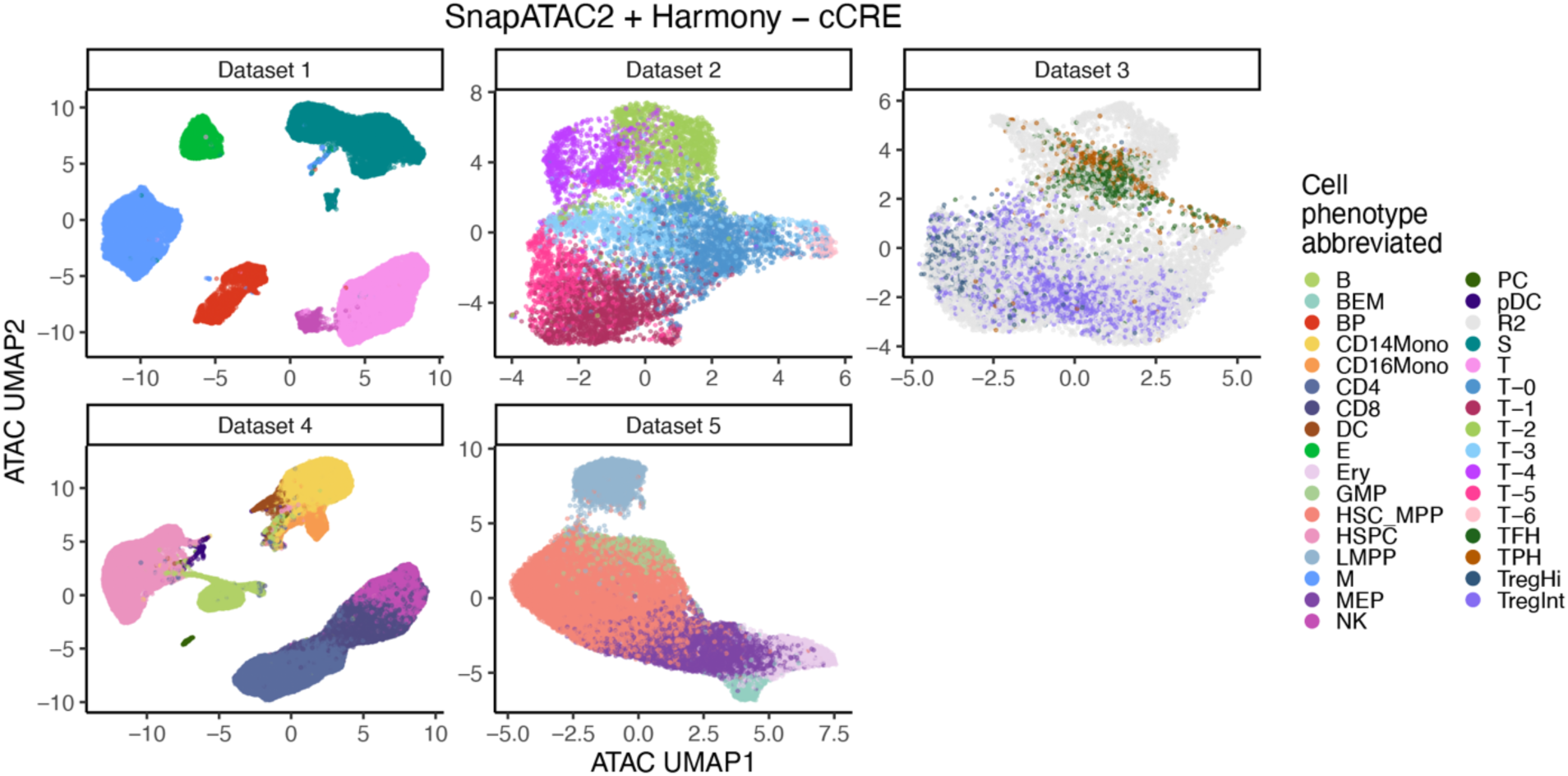
SnapATAC2 preformed best for NN metrics with RNA-embedding1. snATAC-seq UMAPs colored by cell phenotype for all Benchmarking Datasets using SnapATAC2 + Harmony with cCRE features.

**Fig. 5.**
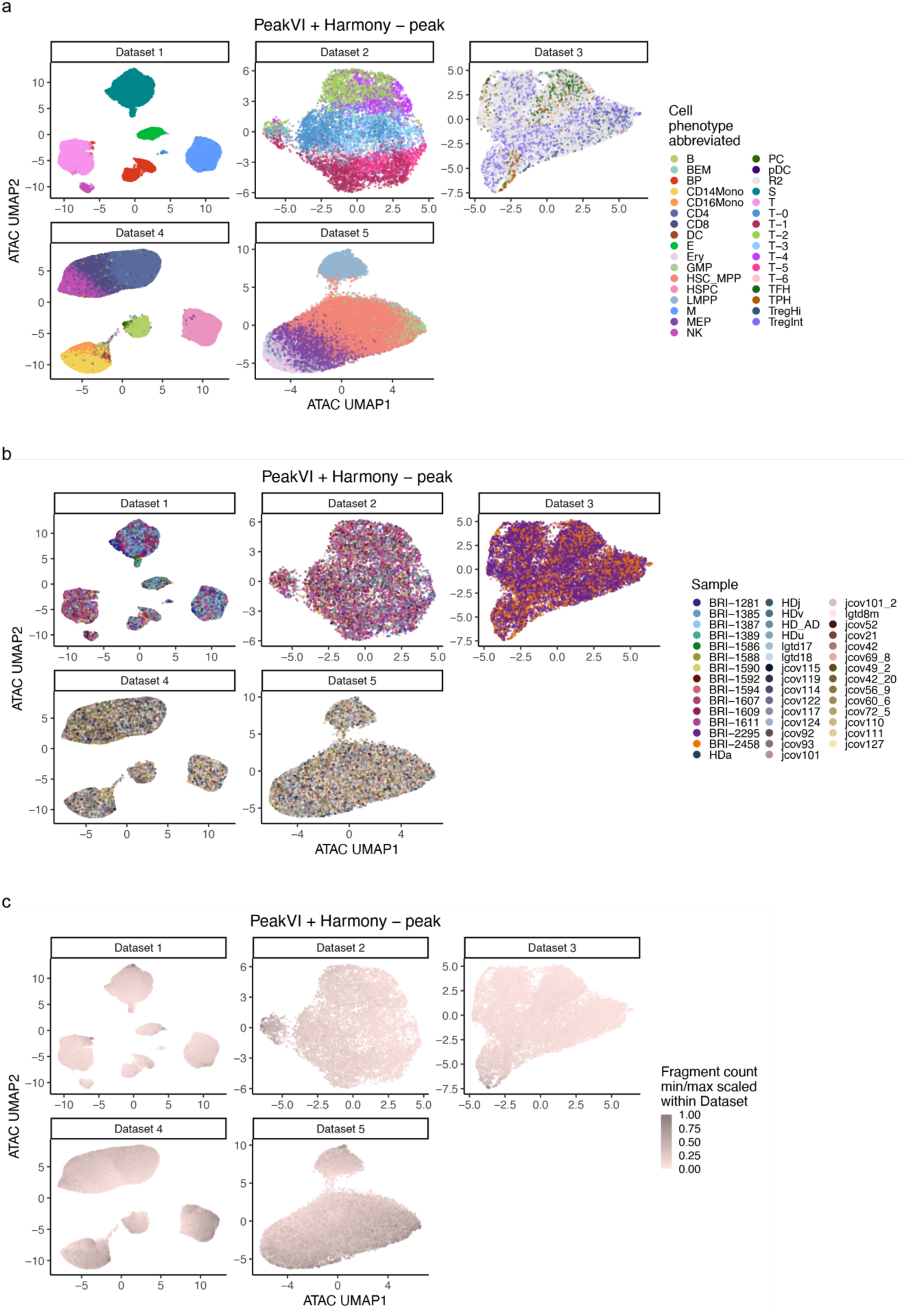
PeakVI biased by fragment count. snATAC-seq UMAPs colored by **a.** cell phenotype, **b.** sample, and **c.** fragment count for all Benchmarking Datasets using PeakVI + Harmony with peak features.

One potential reason that PeakVI performance was suboptimal was confounding by fragment count per cell. We noticed that PeakVI often grouped high fragment count cells together in the same region of the UMAP (**Fig. 5c**). Quantifying this observation, the high fragment counts correlated more with worse bioKNN metrics, corresponding to a worse bio-conservation ranking (**Additional file 2: Fig. S9**). Indeed, in Dataset 3, we saw a group of T cells with higher fragment counts and mixed substates, illustrating poor bio-conservation (**Fig. 5a,c**). This phenomenon was also seen in the pipelines using PeakVI with peak features and PeakVI + Harmony with cCRE features (**Additional file 2: Fig. S10a-b**).

### ArchR itLSI favored cell states over cell types

Using either peaks or cCRE, ArchR’s iterative LSI + Harmony ranked best for Dataset 5 in the RNA-embedding1 NN metrics (score=5, 6.2, respectively; **Fig. 2h**), and demonstrated good performance with the other metrics as well (**Additional file 2: Fig. S7e-f**). They both showed very good separation between the LMPP cluster and the rest on the UMAPs with generally good batch correction (**Additional file 2: Fig. S11a**). itLSI’s best scores were well within the top quartile for the other sub-cell-type resolution datasets: 12.6 for Dataset 2 (peaks with Harmony) and 12.4 for Dataset 3 (tiles with Harmony), out of a theoretical range between 1 and 58 within datasets (**Additional file 2: Fig. S11b**).

However, itLSI performed worse on Dataset 1 and Dataset 4, which had easier to distinguish broad cell types. itLSI + Harmony with cCRE for Dataset 1 was slightly better than the top quartile of scores (score=18.6 vs 19.25) while Dataset 4 using itLSI + Harmony with peaks did not exceed the top quartile (score=24.6 vs 23.85**; Additional file 2: Fig. S11b**). In these datasets, itLSI’s performance was hurt by poor batch effect correction (**Additional file 1: Table S2**). Indeed, Dataset 4’s top pipeline had areas of its UMAPs dominated by single samples, most notably jcov93 within CD4 T & B cells and lgtd18 within T/NK & HSPC cells (**Additional file 2: Fig. S11c**, **left**). Interestingly, the un-Harmonized version scored similarly (score=25.2) with both pipelines having inappropriate mixing of the myeloid, HSPC, B, and pDC cell types in the late COVID-19 samples (**Additional file 2: Fig. S11c**).

### CellSpace performed poorly across datasets, features, correction, and metrics

CellSpace was the worst-ranked integration method across all metrics (**Fig. 2f**; **Additional file 2: Fig. S7a-b**). The best RNA-embedding1 overall ranking CellSpace achieved was for Dataset 4, peaks, without Harmony (score=32.2; **Additional file 1: Table S2**). However, even with distinct cell types in Dataset 4, CellSpace remained as a singular entity in the UMAP, with intermixed domains for CD4 T + CD8 T + NK cells, B + Plasma cells, CD14 Monocytes + CD16 Monocytes + Dendritic cells, HSPC + plasmacytoid dendritic cells (**Additional file 2: Fig. S12a**). However, it was very well integrated at the sample level even without Harmony correction (**Additional file 2: Fig. S12b**), suggesting an over-integration to the point of unclear cell types.

### Harmony improves batch correction for most methods

As Harmony was created to correct batch structure while maintaining biological identity, we expected that it would improve batch correction rankings while minimally affecting bio-conservation rankings (**Fig. 2g**). Indeed, we observed that using Harmony generally improved most methods (**Fig. 6a; Additional file 2: Fig. S13**). The methods that did not inherently address batch effects, PCA, LSI, SnapATAC2, generally benefitted the most. rLSI, rCCA, and PeakVI, which all explicitly corrected for batch, had a more modest benefit with Harmony. The two methods with some implicit batch correction, CellSpace and itLSI, were the most likely to have adverse effects post-Harmony. CellSpace’s poor interaction with Harmony was seen mostly clearly in cell state Datasets 2 and 5, across all ATAC features, with peaks as the most extreme (**Fig. 6a**). However, we saw batch effects for CellSpace both with and without Harmony for Dataset 2 peaks (**Fig. 6b**). Iterative LSI was primarily negatively affected by Harmony in the cell type datasets for GAS features (**Fig. 6a**). This was typified by Dataset 1 when using itLSI + Harmony with basicGAS features, where stronger batch effects were seen in the post-Harmony UMAPs (**Fig. 6c**). However, when itLSI was used with the non-GAS features, including Harmony resulted in similar or better batch correction (**Fig. 6a**).

**Fig. 6.**
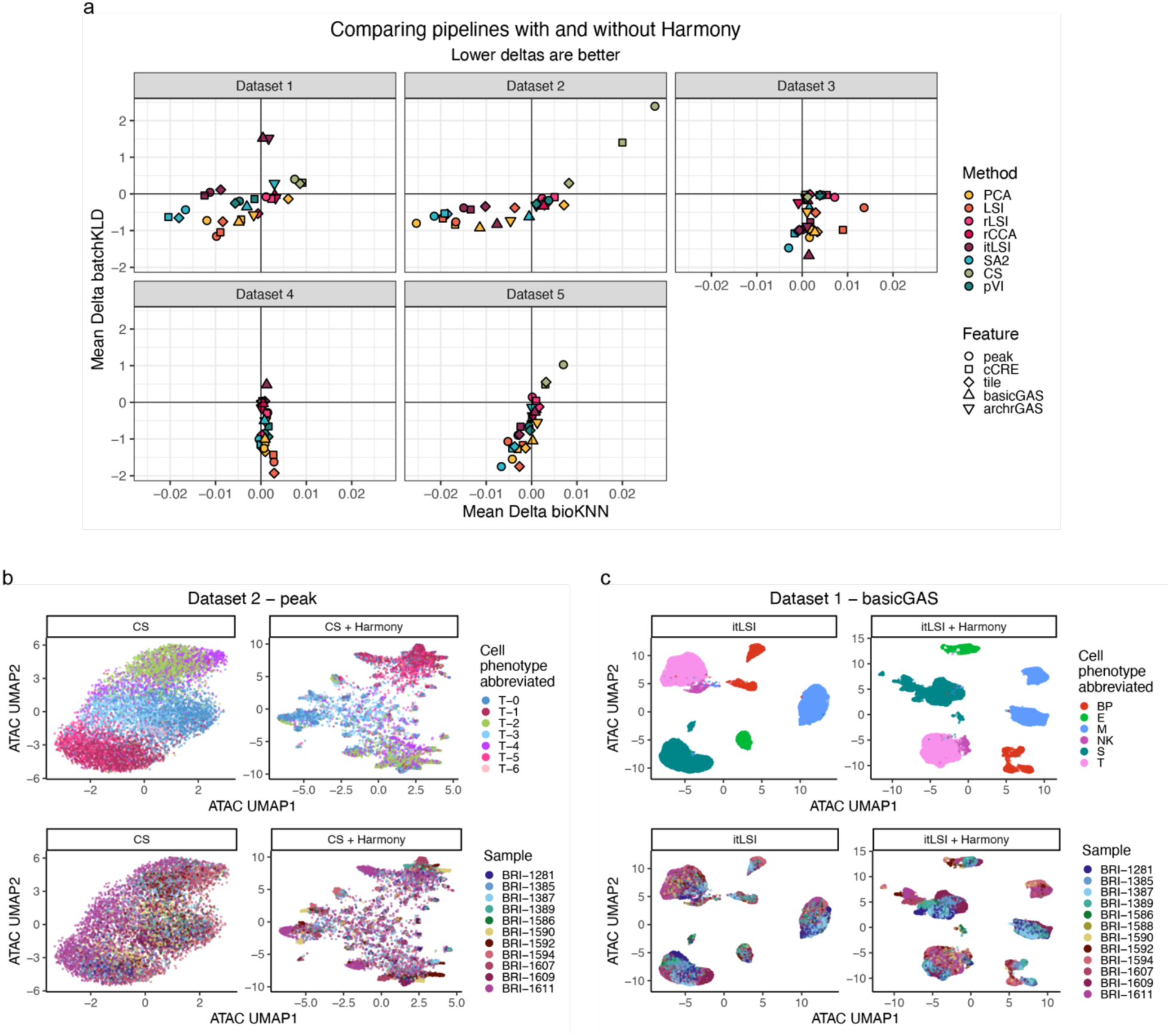
Harmony generally improves batch correction. **a.** Mean bio-conservation (**x-axis**; bioKNN) and batch correction (**y-axis**; batchKLD) Harmony deltas averaged across method/feature combinations in all Benchmarking Datasets. Harmony deltas were calculated as the Harmony metrics subtracted by the no Harmony metrics for the same method/feature combination. Lower deltas were better. **b.** snATAC-seq UMAPs colored by cell phenotype (**top**) and sample (**bottom**) for Dataset 2 with peak features using CellSpace (**left**) or CellSpace + Harmony (**right**). **c.** snATAC-seq UMAPs colored by cell phenotype (**top**) and sample (**bottom**) for Dataset 1 with basicGAS features using ArchR itLSI (**left**) or ArchR itLSI + Harmony (**right**).

### Job requirements varied greatly by method

We assessed time and memory requirements for both the overall pipeline job as well as the individual steps spanning pre-processing, embedding generation, Harmony correction, post-processing, and metric/visualization calculation (**Methods**; **Fig. 7a**; **Additional file 2: Fig. S14-15**). For both time and memory, we found more variation across methods than by input feature. One of the most resource intensive pipelines was rLSI with tiles, requiring 121.6 CPU-hours and 545.7 GB for the ∼200K cells of Dataset 4. PeakVI and CellSpace, two nonlinear methods, also required more time to complete than most. In contrast, the third nonlinear method tested, SnapATAC2, was among the most time and memory efficient. LSI, rCCA, and itLSI were also among the fastest. All methods, excluding rLSI, had roughly similar memory requirements. The use of tiles increased memory usage, likely due to the order of magnitude more features than peaks, cCREs, or GASs. The feature selection step we implemented with CellSpace and PCA (**Methods**) took more memory and often more time than generating the embedding, so a more straightforward feature count cutoff as those used in some of the other methods would decrease the overall job requirements (**Additional file 2: Fig. S14-15**). Also, the metric/visualization steps for Dataset 4 were of high memory and time burden (**Additional file 2: Fig. S14-15**), with the former not necessary to most end-users. Of note, since ArchR required a project to be created before our pipeline could be implemented within it, it was not included in the aggregated plot. It maxed out at 617.2 CPU-min and 82.8 GB for Dataset 4 (**Fig. 7b**), thus adding a small, but not negligible, addition to itLSI’s requirements. In all cases, Harmony correction was extremely efficient with a minimal impact on both time and memory requirements.

**Fig. 7.**
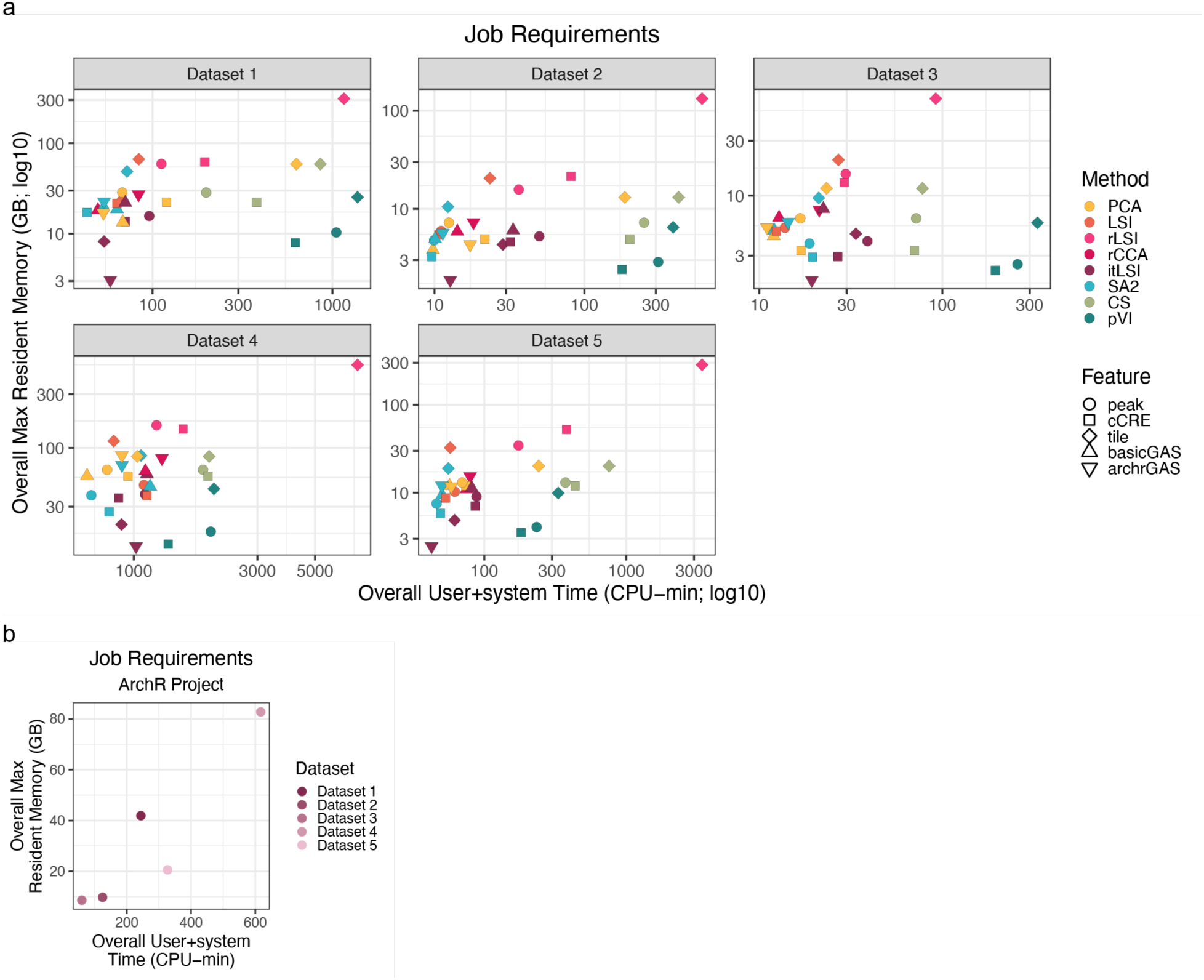
Time and memory requirements vary by method. Time (**x-axis**; user+system time) and memory (**y-axis**; max resident memory) requirements for **a.** entire pipelines and **b.** ArchR project creation for all Benchmarking Datasets. Values calculated with /usr/bin/time (**Methods**).

### NN metrics better for rarer cell phenotypes dominated by fewer samples

The two RNA embeddings used with the NN metrics produced very concordant rankings, with a mean bioKNN correlation of 0.97 and a mean batchKLD correlation of 0.91 (**Additional file 2: Fig. S16a-b**). The only notable slight deviation was in Dataset 3 batchKLD metrics, where RNA-embedding2, generated using the rCCA method, favored rCCA/rLSI snATAC-seq embeddings. This likely explains the rLSI discrepancy found with the combined linear model (**Fig. 2f**; **Additional file 2: Fig. S7a-b**).

The RNA-embedding1 NN metrics and LISI metrics were generally concordant and prioritized pipelines in a similar fashion (**Additional file 2: Fig. S17a-b**). Mean bio-conversation metric correlation across datasets was 0.92; batch correction metrics were slightly less correlated at a mean of 0.75.

However, there were a few discrepancies. The most dissimilar pipeline in the ranked batch correction metrics was rCCA with GAS features, with or without Harmony (**Additional file 2: Fig. S17b**). These pipelines performed well using iLISI, but demonstrated worse performance when assessed with batchKLD. We note that these pipelines were also among the worse-performers in bio-conservation metrics (**Additional file 2: Fig. S17a**), as exemplified by Dataset 5 rCCA with archrGAS features with indistinguishable HSPC states in addition to the very well-mixed samples (**Additional file 2: Fig. S17c-d**). This suggested that these batch-aware rCCA pipelines were likely over-integrating towards iLISI’s uniform sample mixing beyond the local sample distribution expected by the gold standard snRNA-seq embedding, measured by batchKLD.

Furthermore, we saw some interesting examples of cells with good batchKLD values and poor iLISI values, as in the SnapATAC2 + Harmony with cCREs pipeline (**Additional file 2: Fig. S18a**). The most notable disparity was in Dataset 4 (**Additional file 2: Fig. S18b**), where the plasma cells were primarily derived from a single sample (**Additional file 2: Fig. S18c**), seen in the UMAPs for both ATAC (**Fig. 4a**; **Additional file 2: Fig. S18d**) and RNA (**Fig. 1g**; **Additional file 2: Fig. S4c**). In this case of confounded batch and biology, a good result should not have uniform batch mixing, meaning iLISI was an imperfect measure. However, since the gold standard RNA embedding also grouped plasma cells despite belonging to primarily one sample, the sample distributions of the plasma cells’ RNA NN were similar to those of the ATAC NN, as signified by a good batchKLD metric. We saw a similar effect with erythroid progenitors in Dataset 5 (**Additional file 2: Fig. S18e-f**). Therefore, we concluded that batchKLD served as a better metric in cases with rarer cell types that are dominated by fewer samples, which can be where the most interesting biology resides.

## Discussion

In this study, we extensively benchmarked 58 snATAC-seq integration pipelines across 5 datasets, 5 features, 7 methods, 1 embedding correction, and 4 metrics. The performance analysis of these pipelines gave insights into the most effective strategies for single cell chromatin accessibility integration. Based on this, we recommend using ATAC features, such as peaks and cCREs, as opposed to GAS features, which force chromatin accessibility data to look like genes. Furthermore, using Harmony a batch correction step was usually very helpful. We found SnapATAC2 to be the best-performing method in general, though other methods came close in performance, such as PeakVI and ArchR’s itLSI. PCA and LSI were standard methods that performed reasonably well in our benchmark. Our overall best-performing pipeline was SnapATAC2 + Harmony with cCRE, which we recommend for most purposes. In addition to high cross-sample data integration, it had limited time and memory investments, and cCREs render this pipeline to be easily generalizable.

Our study offers a powerful and effective benchmarking strategy. By utilizing multimodal datasets, we could use snRNA-seq embeddings as a standard by which to benchmark snATAC-seq integration. To this end, we introduced two new metrics. We assessed bio-conservation by comparing barcodes for the same cells across modalities in our bioKNN metric; this approach negated the need for potentially imprecise per-cell phenotypic annotations usually defined by functional clusters with hard borders. We measured batch correction using batchKLD, which compared per-cell local sample distributions between modalities and could account for instances where biological states were not uniformly mixed across samples. We note that while RNA embeddings are generally considered to represent key biological features, they can themselves be variable. Thus, we tested two independent embeddings built with two popular approaches within our NN metrics. Our conclusions were generally consistent across RNA embedding choices (**Additional file 2: Fig. S16**). Additionally, we acknowledge that RNA embeddings may be imperfect, and that if there is shared technical variation across modalities, our strategy would not account for it. However, our metrics were also largely consistent with more standard local metrics, cLISI and iLISI (**Additional file 2: Fig. S17a-b**). In certain instances, those metrics may lead to erroneous conclusions where biological states and technical batches co-occur together. For example, the plasma cells in Dataset 4 largely originated from one sample, which iLISI penalized for insufficient mixing, while lsKLD did not as it was validated in the snRNA-seq data (**Additional file 2: Fig. S18b-d**).

Computational analysis of chromatin accessibility data has a fundamental challenge: there is no consistent, reproducible feature set used within and across datasets. Given a new dataset, users often make many specific methodological and parameter choices to call peaks. Reads that do not overlap the final peak set are often discarded, making it possible that the downstream biological conclusions are sensitive to these choices. In terms of data integration, peaks called in one dataset are not easily transferrable across datasets; hence, peaks often need to be recalled and datasets reprocessed. A key finding of this study was the discovery that peaks and ENCODE cCREs achieved similar performance (**Fig. 2f**, **3b**). This suggested that dataset-specific peaks offer little integration advantage over the dataset-agnostic cCREs.

Notably, even though there were no ENCODE DNase-seq datasets assayed with RA synovial tissue or COVID-19 blood, the cCREs retained enough information to identify and integrate our RA and COVID-19 Benchmarking Datasets. Based on this observation, we propose that these pre-defined ENCODE cCREs could be used as a gene-like reference set for future snATAC-seq studies. This would enable easy and reproducible integration with a common feature set, rather than defining bespoke feature sets for each unique application. Furthermore, the deep characterization ENCODE has done of these cCREs provides a rich resource for understanding the functional implications of open chromatin at that locus within the individual datasets. Similar to gene annotation versions, the cCRE annotations could be periodically updated with additional functional elements as datasets continue to increase. We note the possibility that there may be specific biological instances where dataset-specific peaks capture biology missed by ENCODE cCREs.

While we saw general trends across datasets, there were some dataset-specific caveats when selecting appropriate pipelines. For example, datasets are routinely becoming even larger than the 200K cells in Dataset 4; therefore, resource-intensive pipelines like rLSI may become intractable for such datasets. Datasets with uneven fragment counts may lead to biased PeakVI embeddings, particularly for the finer-grain cell substates whose cell phenotype variation is smaller. Those cell state and substate datasets may prefer itLSI while broad cell type datasets may want to avoid it.

We implemented our benchmarking strategy in an easy-to-use command line tool that can be adapted to any dataset of interest, enabling easy benchmarking or inference. After generating the input matrices with the provided feature files and scripts, a user only needs to specify the dataset, feature, and method combinations to get a file of commands that can be run locally or via a computing cluster. All the embeddings, NN graphs, metrics, and UMAPs discussed here were generated in this way.

Our study has some limitations. First, we assume that the underlying cell phenotype will be the same assayed via snRNA-seq and snATAC-seq. In our experience^21^, we have generally observed that cell states that are similar in transcriptional profiles also capture similar biology in chromatin profiles. However, there may be important biological functions captured in one modality, but missed by the other. For example, cell cycle usually affects gene expression more dramatically than chromatin accessibility^30^. Second, our conclusions are based on a limited number of datasets. We chose these datasets to span cell types, states, and substates across both blood and tissue, and we generally observed consistent results across them. However, there were situations where specific pipelines performed slightly better or worse, as discussed above. We also restricted our analysis to the 10x multiome^TM^ platform as it is among the most popular and commercially available, but it is possible that alternative platforms like sci-CAR^31^ or SHARE-seq^30^ may have different results. For the interested reader, we provide the code to apply our framework to other contexts and datasets. Third, we note that our study focused on integration performance. We were unable to quantify factors like software installation, data structure preparation, and ease-of-use per method in an objective fashion. These requirements varied dramatically across pipelines and may be an important factor in user choice.

We hope this benchmarking study will assist researchers decide which combination of feature, method, and correction they will apply to their future snATAC-seq datasets.

## Conclusions

In conclusion, we benchmarked 58 snATAC-seq integration pipelines across 5 datasets utilizing 2 novel multimodal-guided metrics. Using our command-line tool to process these benchmarks, we determined that SnapATAC2 + Harmony using cCRE features outperformed other pipelines while also being generalizable and resource-efficient.

## Methods

### Benchmarking Metrics

#### Bio-conservation K Nearest Neighbors (bioKNN)

To define a gold standard nearest-neighbor graph, we used the snRNA-seq sample-harmonized^20^ PCs (RNA-embedding1) as described in the **Benchmarking Datasets** section. We also used Seurat^29^ version 5 (RNA-embedding2) to test our strategy’s sensitivity to snRNA-seq embedding choice. We used Seurat functions: CreateSeuratObject, split by sample, NormalizeData, FindVariableFeatures, ScaleData, RunPCA, and IntegrateLayers with method=CCAIntegration.

In both cases, we calculated the NN graph from the resulting embedding matrix using RANN::nn2 with K=50, 100, 200 NN. To evaluate each snATAC-seq integration strategy (see **Integration Methods** section), we calculated the K=50, 100, 200 NN as done with the snRNA-seq modality and evaluated what fraction (𝑏𝑖𝑜𝐾𝑁𝑁) of the total NN tested (𝐾) were not represented in both modalities for each cell (𝑁𝑁_𝑠ℎ𝑎𝑟𝑒𝑑_). We wanted lower values to be desirable as in the batchKLD metric.

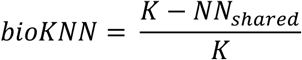

#### Batch Kullback-Leibler Divergence (batchKLD)

To determine how well the samples integrated in the batch-corrected snATAC-seq NN graph, we compared the sample (𝑥) distribution in the ATAC top K NN per cell (𝑄) to that of the gold standard batch-corrected snRNA-seq top K NN for the same cell (𝑃) using KL divergence (**Fig. 1d-e**).

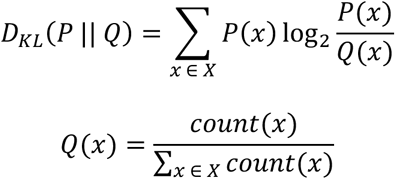

To avoid dividing by 0 and to ensure 𝐾𝐿 = 0 when 𝑃 = 𝑄, we added a pseudocount to 𝑃 and 𝑄. Instead of a uniform pseudocount, we added a dataset-driven pseudocount that accounted for the overall sample distribution of all the cells (𝐴) per dataset, scaled to the equivalent of one additional cell.

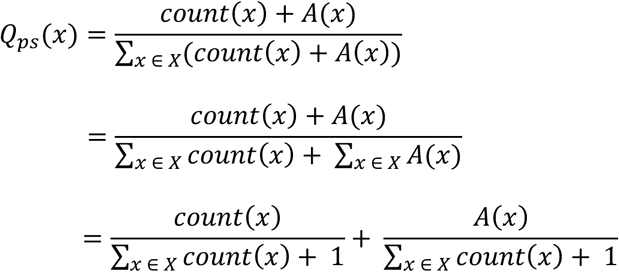

We used the top K NN graphs as described in the *bioKNN* section. We used philentropy::KL to calculate KL divergence; with the addition of the dataset-driven pseudocount, we did not require the epsilon condition of the method, where the small epsilon value was added to 0s in the 𝑄 distribution.

*LISI.* Local Inverse Simpson Index (LISI) scores measured the effective number of different categories of a covariate (𝐶) represented in the local neighborhood of each cell (𝑝_𝑖_, proportional abundances).

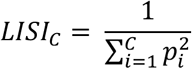

We calculated the LISI scores via lisi::compute_lisi for each embedding matrix for both cell phenotype (cLISI) and sample (iLISI). For all non-substate datasets, the cell type was defined using the multimodal snRNA-seq as described in the **Benchmarking Datasets** section. For the substate dataset (*Dataset 3*), we calculated the cLISI for only the run that had FACS-defined cell types by subsetting both the embedding and metadata matrices to those cells before calculations.

##### ATAC Feature Sets and Matrices

We tested four different genomic feature types: (1) peaks, (2) tiles, (3) cCRE, and (4) two different GAS (**Fig. 1b**; **Table 1**; **Additional file 1: Table S1**).

#### Dataset-specific Peaks

We called peaks from each dataset’s cells as we described previously^21^. For each RA sample, we subsetted the BAM files to chromosomes 1-22XY and QCed cells within each dataset (*Benchmarking Datasets 1-3*) and then converted the subsequent files to MACS2 BEDPE files. For each COVID-19 sample, we started from fragment files requested from the authors, subsetted to the BED-3 columns, chromosomes 1-22XY, and QCed cells from each dataset (*Benchmarking Datasets 4-5*) to get BEDPE files. For each dataset, we concatenated all BEDPE files across samples and called consensus peaks using macs2 callpeak --call-summits with a control file where ATAC-seq was done on free DNA^32^ to account for Tn5’s inherent cutting bias. We trimmed peaks to 500bp (summit +/- 250 bp) and removed overlapping peaks iteratively, keeping the peak with the best q-value. These peaks were overlapped with cell fragments via GenomicRanges::findOverlaps to get a peaks by cells matrix.

#### Tiles

Tiles of 500bp were computed within ArchR^24^ version 1.0.2 via createArrowFiles with addTileMat = TRUE and exported via getMatrixFromProject. The mitochondrial and Y chromosomes were excluded. To calculate fragment overlap, we used GenomicRanges::findOverlaps between ArchR’s tiles and the whole fragments, as done in the other feature types.

#### cCRE

ENCODE SCREEN v3 candidate cis-regulatory elements (cCREs) were downloaded from https://screen.wenglab.org/index/cversions. We also downloaded a set of cell-type-agnostic cCRE with their max DNase-seq z-score from UCSC Data Browser (https://hgdownload.soe.ucsc.edu/gbdb/hg38/encode3/ccre/encodeCcreCombined.bb). For each cCRE, we extended it to 500 bp (midpoint +/- 250 bp) and removed overlapping cCREs iteratively, keeping the cCRE with the highest cell-type-agnostic max DNase-seq z-score. If the cCREs were not cell-type-agnostic or had the same max z-score, then we prioritized by cCRE annotation in the following order: PLS, PLS/CTCF-bound, pELS, pELS/CTCF-bound, dELS, dELS/CTCF-bound, DNase-H3K4me3, DNase-H3K4me3/CTCF-bound, CTCF-only/CTCF-bound. We overlapped the final set of non-overlapping 500bp cCREs with each Benchmarking Dataset’s cell fragments to get a matrix.

#### Gene Activity Scores

We used two gene activity score (GAS) methods. The first was a basic model (‘basicGAS’) that overlapped cell fragments with gene bodies with a 2kb promoter region upstream of the gene. The second was an exponential decay model within ArchR^24^ (‘archrGAS’), calculated via createArrowFiles with addGeneScoreMat = TRUE and exported via getMatrixFromProject. It was then converted from a SummarizedExperiment to a named Sparse Matrix for streamlined processing. In both cases, we used ArchR’s default hg38 gene annotation obtained via getGenes.

##### Integration Methods

We used the above feature matrices as input to a variety of algorithms spanning linear and nonlinear methodologies (**Table 2**). All methods inputted these feature matrices, except ArchR, which reconstructed the feature matrix internally from a user-defined feature set. All non-GAS matrices used 500 bp features; they were also binarized, except for Signac and SnapATAC2 as they specified counts. GAS matrices were processed as if they were gene expression data. Note that not all snATAC-seq integration methods were suited for GAS feature types; if excluded, a justification was given per method. We used defaults for all methods unless otherwise stated. In total, we tested 58 strategies of feature types, integration methods, and Harmony correction (**Additional file 1: Table S1**). For visualization purposes, we calculated a UMAP via umap::umap for each embedding.

*Principal Component Analysis (PCA).* PCA defined orthogonal principal components (PCs) that explained progressively smaller percentages of variation. It is widely used in genomics to reduce data dimensionality, including for single-cell data^33^. We used the snATAC-seq normalization strategy of term frequency-inverse document frequency (TF-IDF) as part of the preprocessing.

We generated PC embeddings for each non-GAS feature type binary matrix via: subsetting features by those with accessibility in at least 0.5% of cells, log(TFxIDF) normalization, variable feature selection, center/scale features, and PCA via irlba::prcomp_irlba^34^. For the basic GAS features, we log-normalized features; ArchR GAS features were already normalized to 10,000, so we logged its matrix with log1p. In both GAS types, we then did variable feature selection, center/scale features, and PCA via irlba::prcomp_irlba. We used 30 dimensions here since the default 3 will likely be too small for this application and all other methods, except PeakVI, defaulted to 30 dimensions. We used irlba R version 2.3.5.1, a package not specifically tailored to snATAC-seq data analysis or pipelines.

#### Signac

Signac^27^ is the commonly used^4,10,22^ snATAC-seq extension of the Seurat package. It used latent semantic indexing (LSI), a combination of TF-IDF normalization followed by Singular Value Decomposition (SVD). For integration purposes, they recommended reciprocal LSI (rLSI), iteratively projecting each dataset into a shared LSI space.

Using Signac version 1.14.0, we processed the snATAC-seq non-GAS non-binary feature matrices with the LSI pipeline: CreateSeuratObject along with the cell metadata, FindTopFeatures, RunTFIDF, and RunSVD. For rLSI in the non-GAS features, we created the Seurat object as before, but split it by sample into a list of objects using SplitObject, and ran the LSI pipeline per object. After merging objects and joining layers, we ran the LSI pipeline again on the combined object. We then used FindIntegrationAnchors and IntegrateEmbeddings to get the rLSI embedding. We calculated the default 30 dimensions, but only used the 2:30 dimensions as suggested by their tutorial as the first dimension was usually correlated to sequencing depth. For GAS features, we used the related pipeline Seurat version 5.1.0, developed primarily for scRNA-seq. However, since Seurat’s default method is PCA, already tested here, we only tested their anchor integration approach using rCCA as described in the *bioKNN* section. For pre-normalized archrGAS features, we built a Seurat object using CreateAssay5Object with the data layer set to the logged matrix before continuing the pipeline outlined above.

We encountered two errors while running rLSI/rCCA. During the rLSI IntegrateEmbeddings step, we would occasionally get FindWeights errors saying “Number of anchor cells is less than k.weight. Consider lowering k.weight to less than XX or increase k.anchor”, where XX was an integer. We reran the same command with an additional argument k.weight=XX-1 until the error no longer occurred. During rCCA with the pre-normalized archrGAS features, we got FindVariableFeatures errors in match.arg. We reran the same command with selection.method=”mvp” instead of the default “vst”.

#### ArchR

ArchR^24^ is a popular^6,7,35^ end-to-end pipeline for analyzing scATAC-seq data. It used iterative LSI for dimensionality reduction and implicit batch correction. For the first iteration, ArchR used the top accessible features (by default, 25,000 500 bp tiles). Subsequently, it calculated LSI, defined clusters, sum-aggregated accessibility per cluster, log-normalized, and identified most variable features across clusters to use in the next, and by default final, round of LSI. ArchR did not explicitly correct for batch within this process, claiming that the first round of LSI identified low resolution clusters that were not batch confounded. We did not use ArchR’s estimated LSI functionality as it was not set by default.

We inputted the per-sample post cell QC fragment files to ArchR version 1.0.2 createArrowFiles with all additional QC flags nullified to avoid subsetting on cells; the tile and archrGAS matrices were created in this step as mentioned previously. We then created an ArchRProject. Since ArchR does not allow non-gene-expression matrices to be inputted directly, we used the feature sets calculated in **ATAC Feature Sets and Matrices** to build the feature matrices within the ArchRProject. The peak matrix was added to the project via addPeakSet and addPeakMatrix while the cCRE matrix was added via addFeatureMatrix. Since the basicGAS matrix was count data, it was converted to a SummarizedExperiment object and inputted with addGeneExpression to allow for the calculation of gene-specific parameters. For each non-GAS feature matrix, we applied the IterativeLSI process via addIterativeLSI with default parameters.

For the GAS feature matrices, we used addIterativeLSI with parameters to mimic scRNA-seq: binarize = FALSE, firstSelection = "var", varFeatures = 2000. In both cases, we calculated the default 30 dimensions, though of note, ArchR excluded dimensions correlated to sequencing depth by default. ArchR did not require a batch covariate. The embedding matrices were exported from the ArchR project via getReducedDims and converted from a SummarizedExperiment to a named Sparse Matrix for streamlined postprocessing.

#### SnapATAC2

SnapATAC2^23^ used a matrix-free spectral embedding for dimensionality reduction. This nonlinear method was built on the Lanczos algorithm to implicitly use the Laplacian matrix without storing it to decrease time and space complexity. SnapATAC2 utilized independent batch correction algorithms in their paper, with Harmony^20^ among its suggested set.

Since SnapATAC2 was written in python, we first converted the nonbinary R Sparse Matrix to MatrixMarket format using writeMM. We then created an AnnData object using snapatac2.read_10x_mtx with that matrix file, an observation file including cell metadata, and a variable file of peak names. For non-GAS features, we selected 50,000 features with snapatac2.pp.select_features, as suggested in their tutorial; this was the only post-IO step in their pipeline before non-GAS embedding creation. Since basicGAS features are counts, we used scanpy functions as suggested by their paper: scanpy.pp.highly_variable_genes with flavor=’seurat_v3’ before subsetting to those variable genes, scanpy.pp.normalize_total, scanpy.pp.log1p. As the archrGAS features were already normalized, we used scanpy.pp.log1p and scanpy.pp.highly_variable_genes with flavor=’seurat’ before subsetting. We then ran their dimensionality reduction method with snapatac2.tl.spectral with a seed and the default 30 dimensions set and if a GAS feature, features=None. Note that with default weighted_by_sd=TRUE, the resulting matrix could be less than 30 dimensions. We used SnapATAC2 version 2.5.3. We then extracted the embedding matrix to a text file that we loaded back into R to save as a named Sparse Matrix. SnapATAC2 did not require a batch covariate.

*CellSpace.* CellSpace^28^ co-embedded k-mers and cells to infer features and TF motifs from their constituent k-mers using neural embedding modeling software StarSpace^36^. It randomly sampled overlapping k-mers from inputted accessible features, peaks or tiles preferred, to generate training examples of k-mers accessible per cell (“positive” cell). Negative cells without accessibility for the k-mer set were randomly sampled since there were many more inaccessible cells for any given feature; this also helped with false negatives and sparsity. During training, k-mer and cell embeddings were updated to move the induced feature sequence embedding closer to positive cells and further from negative cells. N-grams of flanking sequence around k- mers provided additional context. Batch variables were not used here as the authors claim the final embedding generated from small k-mers was less influenced by batch effects. We did not run the GAS matrices here since randomly sampling 8-mers across an entire gene seemed counterintuitive to the TF-centric theme of this method.

We downloaded and compiled CellSpace version 1.0.0 as their GitHub tutorial recommended. CellSpace required a file of cell names, the fasta file of feature sequences, and a MatrixMarket formatted matrix. CellSpace did not internally do feature selection, though its paper recommended using top variable features to speed up runtime and possibly improve quality.

Therefore, we subsetted to variable features calculated as in the PCA pipeline discussed above before converting from R Sparse Matrix to MatrixMarket using writeMM. We generated fasta files of the feature regions via GenomicRanges::getSeq. We ran CellSpace with all default parameters, except requesting 5 threads instead of 10. CellSpace did not require a batch covariate. The resulting tsv file was then inputted into its corresponding R package CellSpace function to access and save its embedding slot cell.emb as a 30-dimension named Sparse Matrix.

#### PeakVI

As part of the scVI suite of tools, PeakVI^25^ used a variational autoencoder (VAE) to model a latent space conditioned on a user-specified batch variable to correct for those batch effects and capture batch-independent biological variation. The batch variable was also used in the decoding step to calculate the probability of accessibility, which was further modified to the probability of observation by multiplying by the region-specific factors and cell-specific factors calculated from the input. Because PeakVI modeled a Bernoulli distribution, we did not use the non-binary GAS matrices as input.

Since PeakVI was written in python, we first converted the binary R Sparse Matrices to MatrixMarket format using writeMM before converting into a python AnnData format using scvi.data.read_10x_atac. Cell metadata was added to the AnnData object separately, verifying that all observations and variables were unique. We then filtered the features to keep those in at least 5% of cells as recommended by their tutorial. We setup the AnnData object using scvi.data.setup_anndata with batch_key=’sample’ before training the PeakVI model with scvi.model.PEAKVI with all default parameters. We used scvi-tools version 1.1.6. We then extracted the embedding matrix to a text file that we loaded back into R to save as a named Sparse Matrix.

##### Harmony Correction

To standardize the Harmony implementation and defaults, we applied the stand-alone Harmony package version 1.2.1 to each integration method’s output embedding, using all dimensions given. We batch corrected by sample using default parameters.

##### Benchmarking Datasets

An overview of the benchmarking datasets used in this study is in **Table 3**. RNA UMAPs with gold standard biological cell phenotypes defined from RNA or protein can be found in **Fig. 1g**; these annotations were used in the cLISI bio-conservation metric (see **Benchmarking Metrics** section). Additionally, as part of the Seurat NN graphs generated in the **Benchmarking Metrics** section, we removed samples with less than 100 cells from the Benchmarking Datasets before any further processing since Seurat’s anchor integration had issues with very small cell count batches. **Additional file 2: Figs. S1-S5** describe each dataset: panels **a**-**b** denote quality control statistics for each modality, panels **c**-**e** denote snRNA-seq batch correction (iLISI) and bio-conservation (cLISI) metrics, and for the RNA-based cell phenotypes, panels **f**-**i** denote comparisons between our reprocessing and the original paper’s processing. Full methodological details of the initial processing can be found in the original studies^21,22^, with summaries here.

#### Benchmark Dataset 1. RA inflammatory tissue broad cell types

This dataset^21^ consisted of 12 synovial tissue samples, 11 from RA patients and 1 from an osteoarthritis (OA) patient, and included 6 broad cell types (B/plasma, T, NK, myeloid, stromal [fibroblasts/mural], and endothelial) across 31,547 cells (median 3,093 cells/sample). Each sample was processed independently at the cell capture step and run through a 10x multiome experiment. To be included, cells had to pass quality control metrics in both snRNA-seq and snATAC-seq modalities (post-QC in **Additional file 2: Fig. S1a-b**) as well as have the same annotated broad cell type. We reprocessed the snRNA-seq data using the updated software versions uniformly used in this study. Briefly, we normalized, selected variable genes, centered/scaled genes, computed 20 PCs, batch corrected by sample via Harmony^20^, created a shared nearest neighbor graph, clustered with Louvain clustering, and generated a UMAP. This reprocessing generated the RNA-embedding1 used in the NN metrics. It showed good batch correction (**Additional file 2: Fig. S1c-d**) and cell type bio-conservation (**Additional file 2: Fig. S1e**). We chose a clustering resolution that closely mirrored the original processing (**Additional file 2: Fig. S1f-i**).

#### Benchmark Dataset 2. RA inflammatory tissue T cell states

This dataset^21^ used 8,069 T cells from 11 of the RA/OA synovial tissue samples discussed above; one sample was dropped for having less than 100 cells. As the original study did not define snRNA-seq cell states from the multiome data alone, we clustered as above using 10 sample-harmonized^20^ PCs. Samples were well-mixed after Harmony (**Additional file 2: Fig. S2c-d**) with consistent clusters (**Additional file 2: Fig. S2e**). We chose a clustering resolution that corresponded to the chromatin classes defined by the original study (**Additional file 2: Fig. S2f-g**). We annotated cell states using marker genes similar to those in the original study and via differential gene analysis using presto::wilcoxauc on the normalized genes x cells matrix (**Additional file 2: Fig. S2h-i**).

#### Benchmark Dataset 3. RA PBMC T cell substates

This dataset^21^ totaled 10,669 cells across two runs of four pooled RA PBMC samples each. There were no donors shared between runs, and the donors were not genotyped, disallowing genotype-based demultiplexing. After enriching for CD4+ T cells, four populations were sorted using Fluorescence-Activated Cell Sorting (FACS): CD4^+^CD127^-^CD25^hi^ Tregs, CD4^+^CD127^-^CD25^int^ Tregs, CD4^+^CD25^-^PD1^+^CXCR5^+^ TFH, and CD4^+^CD25^-^PD1^+^CXCR5^-^ TPH. Each population was tagged with a hashing antibody. We generated 10x multiome data from these cells with an HTO library generated to label the specific populations. Cells had to pass quality control metrics in all modalities processed.

Overall, the quality was slightly worse than the RA tissue multiome datasets in *Benchmark Datasets 1-2* with fewer fragments and genes detected and higher mitochondrial reads (**Additional file 2: Fig. S3a-b**). We used 10 run-harmonized PCs to generate a snRNA-seq UMAP (**Fig. 1g**) and a gene KNN graph for use in the bioKNN and batchKLD metrics as defined above. Samples mixed reasonably well (**Additional file 2: Fig. S3c-d**), though there was a cell imbalance between runs (2,998 in Run 1 vs 7,671 in Run 2). While both runs were used for all snRNA-seq and snATAC-seq processing, only one run had processed FACS populations, so the cLISI calculation was restricted to that run in both modalities (**Additional file 2: Fig. S3e**).

#### Benchmark Dataset 4. COVID-19 PBMC broad cell types

This study^22^ profiled mature and progenitor cell populations from peripheral blood samples using PBMC-PIE (peripheral blood mononuclear cell analysis with progenitor input enrichment). This 10x multiome dataset was comprised of 30 PBMC samples collected between March 2020 and March 2021 from 26 individuals, including subsets of healthy adults, adults recovering from a non-COVID-19 critical illness, and adults at early (2-4 months) or late (4-12 months) post-infection stages of COVID-19 convalescence. In the original study, sample merging (using Seurat^29^ for snRNA-seq and Signac^27^ for snATAC-seq), quality control, doublet removal (using Scrublet for snRNA-seq and Amulet for snATAC-seq), batch correction by sample with Harmony^20^, and iterative clustering/annotation were preformed to get a final dataset of 197,360 cells (median 6,027 cells/sample). Overall, there were fewer genes and fragments found here with a much higher mitochondrial percentage than in *Benchmark Datasets 1-2* (**Additional file 2: Fig. S4a-b**).

Using marker gene expression, the authors defined 10 major cell type clusters: NK, CD8+ T, CD4+ T, B, plasma, CD16+ monocyte, CD14+ monocyte, DC, pDC, and hematopoietic stem and progenitor cells (HSPC). For uniform processing and to generate a nearest neighbor graph for use in the NN metrics, we reprocessed the multiome snRNA-seq data using the pipeline denoted in *Benchmark Dataset 1* using 30 sample-harmonized PCs. We saw good sample mixing (**Additional file 2: Fig. S4c-d**) and internal cell type consistency (**Additional file 2: Fig. S4e**). Furthermore, we compared the reprocessed cell types to the originally-defined cell types (**Additional file 2: Fig. S4f-i**) and saw general consistency.

#### Benchmark Dataset 5. COVID-19 PBMC HSPC states

In the original paper, the 28,069 HSPCs in the previous dataset ^22^ were further subclustered into 6 states: hematopoietic stem cells and multipotent progenitors (HSC_MPP), lymphoid-primed MPP (LMPP), granulocyte-monocyte progenitors (GMP), megakaryocyte-erythroid progenitors (MEP), erythroid progenitors (Ery), and basophil-eosinophil-mast (BEM) cell progenitors. As above, we dropped 2 samples with fewer than 100 cells and re-annotated cells using 20 multimodal snRNA-seq sample- harmonized PCs (**Additional file 2: Fig. S5c-e**). We concatenated the multiple HSC_MPP clusters as the original authors did. The cluster borders were highlighted in a higher (lighter) cLISI score in **Additional file 2: Fig. S5e**. We compared them to the original marker gene annotations and saw common cell states (**Additional file 2: Fig. S5f-i**).

##### Pipeline

After creating all the feature matrices defined in **ATAC Feature Sets and Matrices** for each dataset defined in **Benchmarking Datasets**, we used a command line tool we developed to create all embeddings, Harmony-corrected embeddings, NN graphs, metrics, and UMAPs.

The datasets and features were inputted into this pipeline via a “dictionary file,” which connected dataset and feature keywords to file paths for the required types of files: feature matrix, features for ArchR input, ArchR project, metadata, and gene NN graphs. These dataset and feature keywords were then referenced in the pipeline script to generate dataset/feature/method-specific command files that we piped into a SLURM scheduler. Therefore, any dataset or feature combination future users desire can be used to generate the types of output for each method we detailed here.

##### Linear modeling

We used two different linear models, combined and separated, to assess the overall metric ranking for each of the three metric types: (1) NN metrics using RNA- embedding1, (2) NN metrics using RNA-embedding2, (3) LISI metrics. The bio-conservation metrics for NN and LISI were bioKNN and cLISI, respectively while the batch metrics were batchKLD and iLISI, respectively. Each metric was mean-averaged across cells per dataset, feature, method, and correction combination and subsequently ranked within dataset where 1 is the best rank and 58 is the worst rank (**Additional file 1: Table S2-S4**). In both models, this rank was used to create a score that was related to the dataset, feature, method, and correction combination using stats::glm in R with family=”gaussian”.

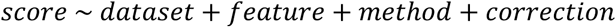

In the combined model, using percentages based off Luecken et al.,^15^, we ran 1 model using

score:

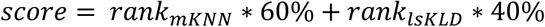

In the separated model, we ran 2 models using scores:

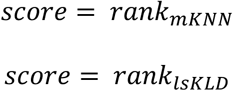

To avoid overdetermination in the model, we chose a reference value for each covariate as follows: dataset – dataset1, feature – tile, method – LSI, and correction – No Harmony. Each covariate’s matrix was 1-hot encoded.

##### Job Requirements

We assessed time and memory requirements per dataset/feature/method pipeline job and per step within a pipeline job using /usr/bin/time. We summed both system and user CPU-minutes for time while using max resident set size (RSS) to assess memory. Steps were separated into 5 categories. Firstly, pre-processing included variable peak/gene selection, data type conversions from R to python, and feature matrix re-creation within ArchR projects.

Secondly, embeddings were generated and thirdly, they were corrected with Harmony. Fourthly, post-processing was converting both uncorrected and corrected embeddings back into R Matrices. Fifthly, NN graphs, metrics, and UMAPS were generated for metrics/visualization purposes for both uncorrected and corrected embeddings.

## Declarations

### Ethics approval and consent to participate

Not applicable

### Consent for publication

Not applicable

### Availability of data and materials

RA datasets analyzed during this study are included in Weinand et al., Nat Commun, 2024^21^ and available from the Synapse repository using identifier syn53650034^37^ (https://doi.org/10.7303/syn53650034). COVID-19 datasets analyzed during this study are included in Cheong et al., Cell, 2023^22^ and available from the GEO repository using identifiers GSE196987 and GSE196988 (https://www.ncbi.nlm.nih.gov/geo/query/acc.cgi?acc=GSE196987; https://www.ncbi.nlm.nih.gov/geo/query/acc.cgi?acc=GSE196988).

Processed data generated in this study are available from the Zenodo repository using identifier 15073137 (https://doi.org/10.5281/zenodo.15073137). A website of UMAPs generated in this study is available at https://immunogenomics.io/snATAC_benchmark/.

The command-line tool code used to generate the results and figures presented in this study can be found on GitHub (https://github.com/immunogenomics/snATAC_benchmark/). We have also listed there the conda and R packages we used.

### Author contributions

KW and SR conceptualized the study. KW defined the methodology, software, and visualization with input from EL and SR. KW processed and analyzed the Benchmarking Datasets. EL curated and initially processed and analyzed the COVID-19 datasets. MC designed the website. KW and SR drafted the original manuscript with edits by EL and MC. SR supervised the study and provided funding. All authors read and approved the final manuscript.

## Acknowledgements

We thank Aparna Nathan and Zepeng Mu for their helpful feedback on this manuscript. We also thank Kamil Slowikowski for website assistance.

## Additional files

Additional file 1. Supplementary Tables S1-S4

Excel Spreadsheet, .xlsx

Additional file 2. Supplementary Figures S1-S17

Word Document, .docx

