## Supplementary Figures for "Defining effective strategies to integrate multi-sample single-nucleus ATAC-seq datasets via a multimodal-guided approach"

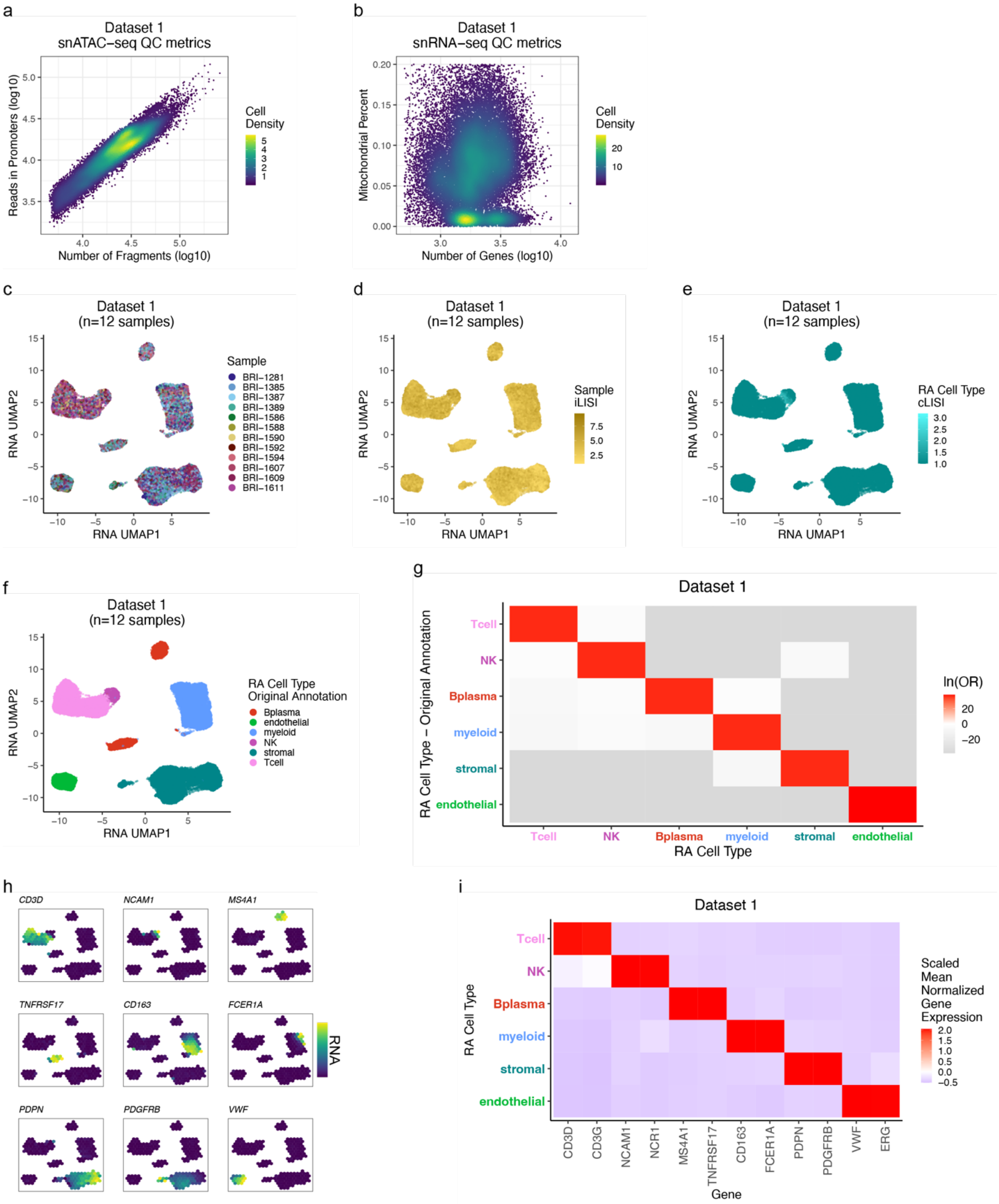


**Fig. S1.** Dataset 1. RA inflammatory tissue broad cell types.
**a.** snATAC-seq QC metrics of fragment count and reads in promoters, colored by cell density.
**b.** snRNA-seq QC metrics of gene count and mitochondrial percent, colored by cell density.
**c.** snRNA-seq UMAP colored by sample.
**d.** snRNA-seq UMAP colored by sample integration LISI (iLISI). Higher values (darker colors) are indictive of better batch correction.

**e.** snRNA-seq UMAP colored by cell type LISI (cLISI). Lower values (darker colors) are indictive of better bio-conservation.
**f.** snRNA-seq UMAP colored by broad cell types defined in the original study[21].
**g.** Natural log of the odds ratio between the re-annotated and original broad cell types from **f**. Non-significant values (FDR>0.05) are white.
**h.** Binned mean-normalized marker gene expression on snRNA-seq UMAP; yellow denotes high expression.
**i**. Scaled mean-normalized marker gene expression across re-annotated broad cell types.

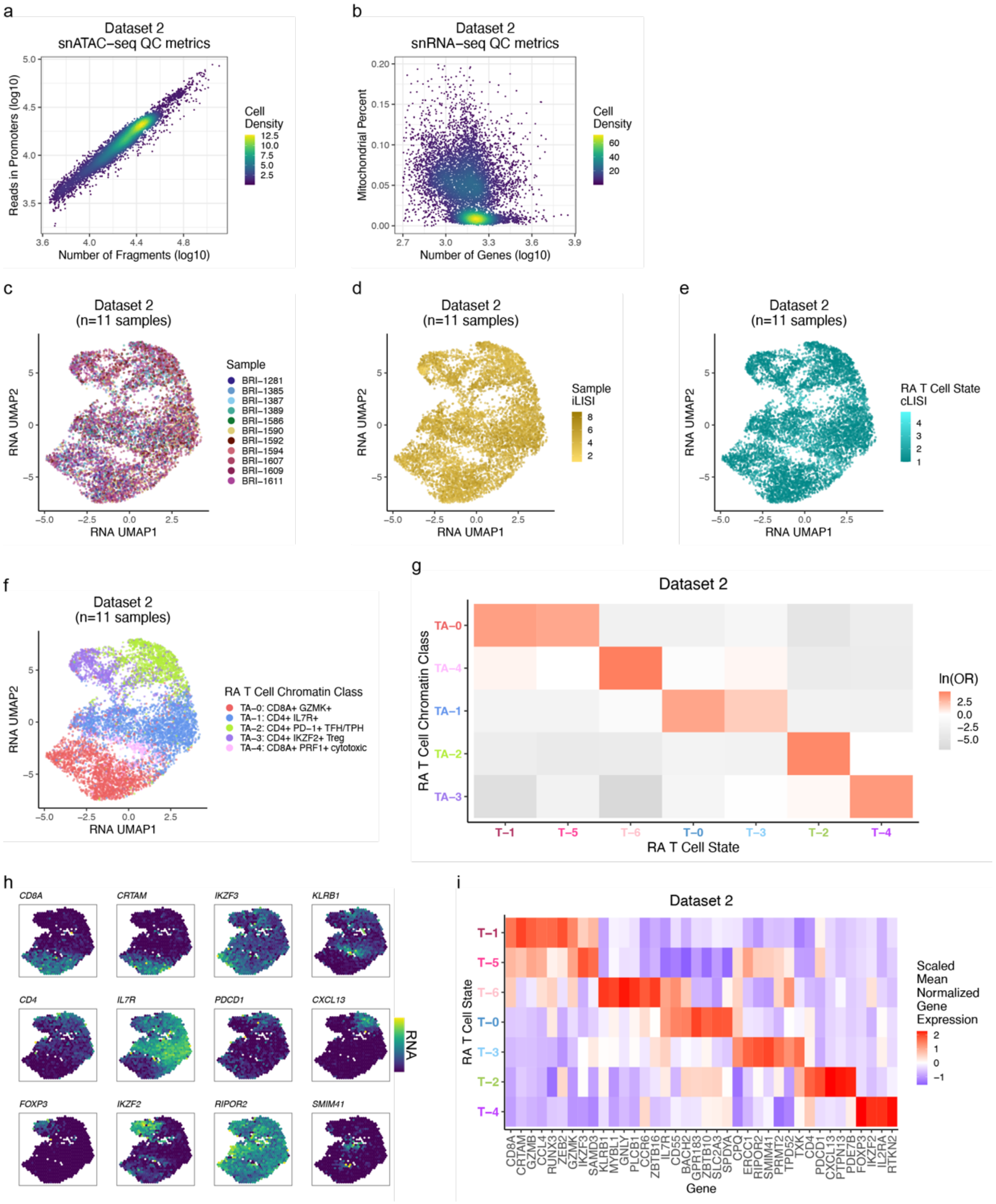


**Fig. S2.** Dataset 2. RA inflammatory tissue T cell states.
**a.** snATAC-seq QC metrics of fragment count and reads in promoters, colored by cell density.
**b.** snRNA-seq QC metrics of gene count and mitochondrial percent, colored by cell density.
**c.** snRNA-seq UMAP colored by sample.
**d.** snRNA-seq UMAP colored by sample integration LISI (iLISI). Higher values (darker colors) are indictive of better batch correction.
**e.** snRNA-seq UMAP colored by cell type LISI (cLISI). Lower values (darker colors) are indictive of better bio-conservation.
**f.** snRNA-seq UMAP colored by RA T cell chromatin classes defined in the original study[21].
**g.** Natural log of the odds ratio between the RNA-defined RA T cell states and the chromatin classes from **f**. Non-significant values (FDR>0.05) are white.

**h.** Binned mean-normalized marker gene expression on snRNA-seq UMAP; yellow denotes high expression.
**i**. Scaled mean-normalized marker gene expression across RNA-defined RA T cell states.

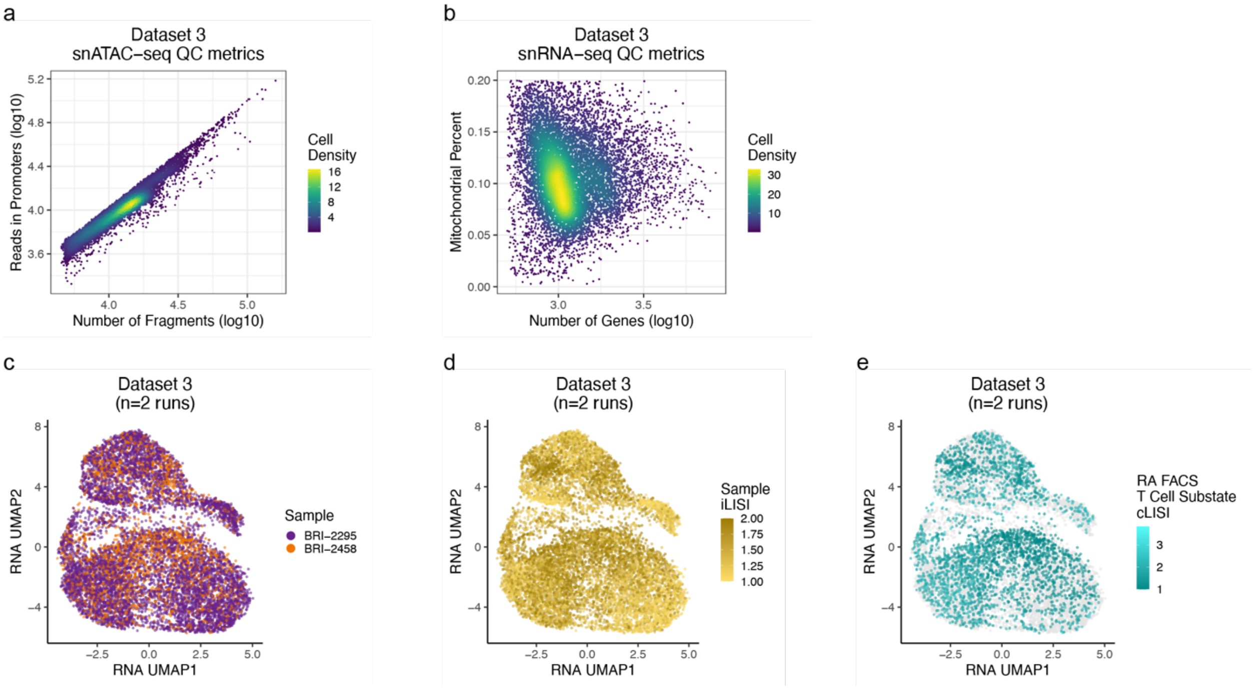


**Fig. S3.** Dataset 3. RA PBMC T cell substates.
**a.** snATAC-seq QC metrics of fragment count and reads in promoters, colored by cell density.
**b.** snRNA-seq QC metrics of gene count and mitochondrial percent, colored by cell density.
**c.** snRNA-seq UMAP colored by sample.
**d.** snRNA-seq UMAP colored by sample integration LISI (iLISI). Higher values (darker colors) are indictive of better batch correction.
**e.** snRNA-seq UMAP colored by cell type LISI (cLISI). Lower values (darker colors) are indictive of better bio-conservation. Light grey values are from Run 2 that did not have FACS labels.

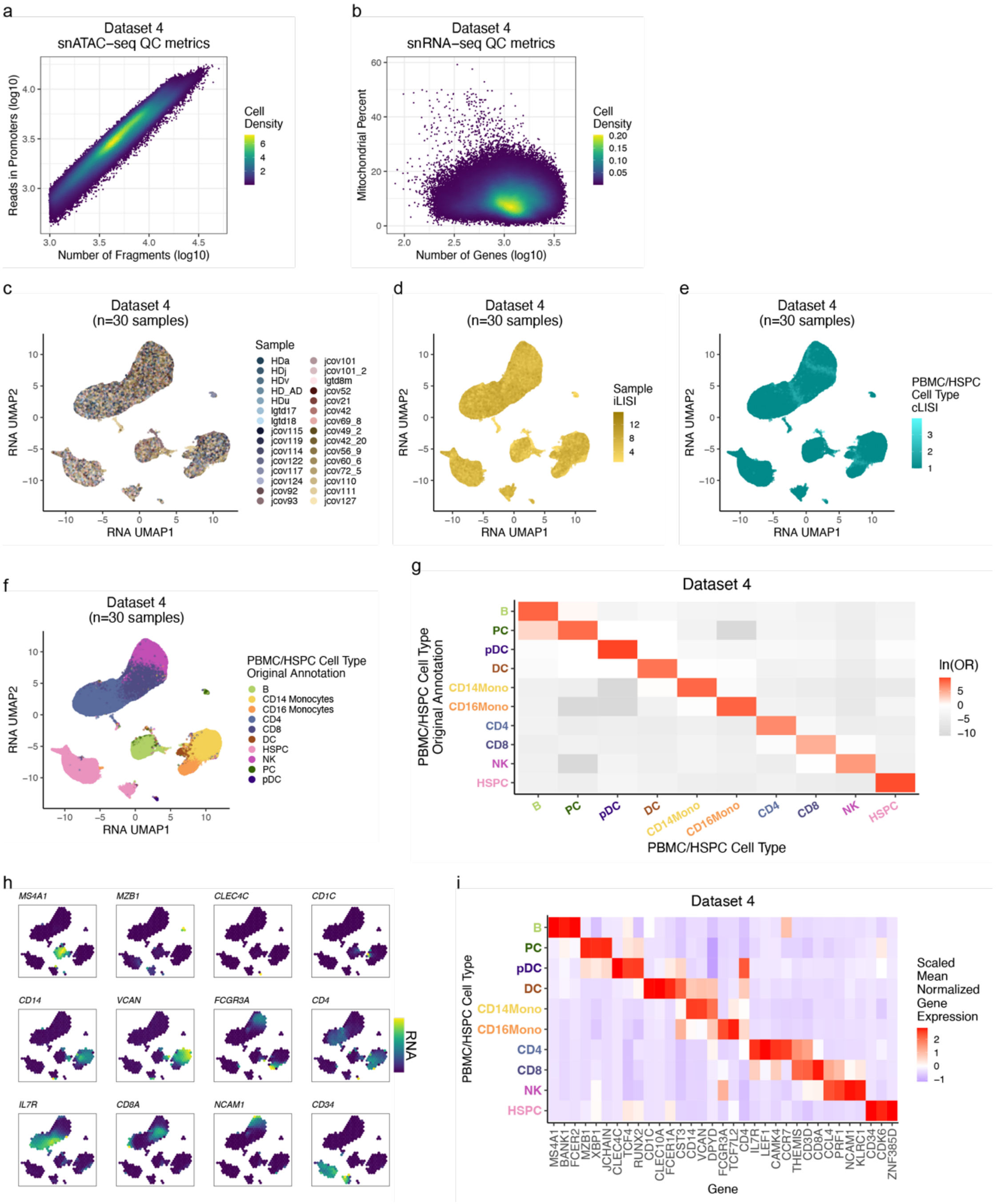


**Fig. S4.** Dataset 4. COVID-19 PBMC broad cell types.
**a.** snATAC-seq QC metrics of fragment count and reads in promoters, colored by cell density.
**b.** snRNA-seq QC metrics of gene count and mitochondrial percent, colored by cell density.
**c.** snRNA-seq UMAP colored by sample.
**d.** snRNA-seq UMAP colored by sample integration LISI (iLISI). Higher values (darker colors) are indictive of better batch correction.
**e.** snRNA-seq UMAP colored by cell type LISI (cLISI). Lower values (darker colors) are indictive of better bio-conservation.
**f.** snRNA-seq UMAP colored by broad cell types defined in the original study[22].
**g.** Natural log of the odds ratio between the re-annotated and original broad cell types from **f**.

Non-significant values (FDR>0.05) are white.

**h.** Binned mean-normalized marker gene expression on snRNA-seq UMAP; yellow denotes high expression.

**i**. Scaled mean-normalized marker gene expression across re-annotated broad cell types.

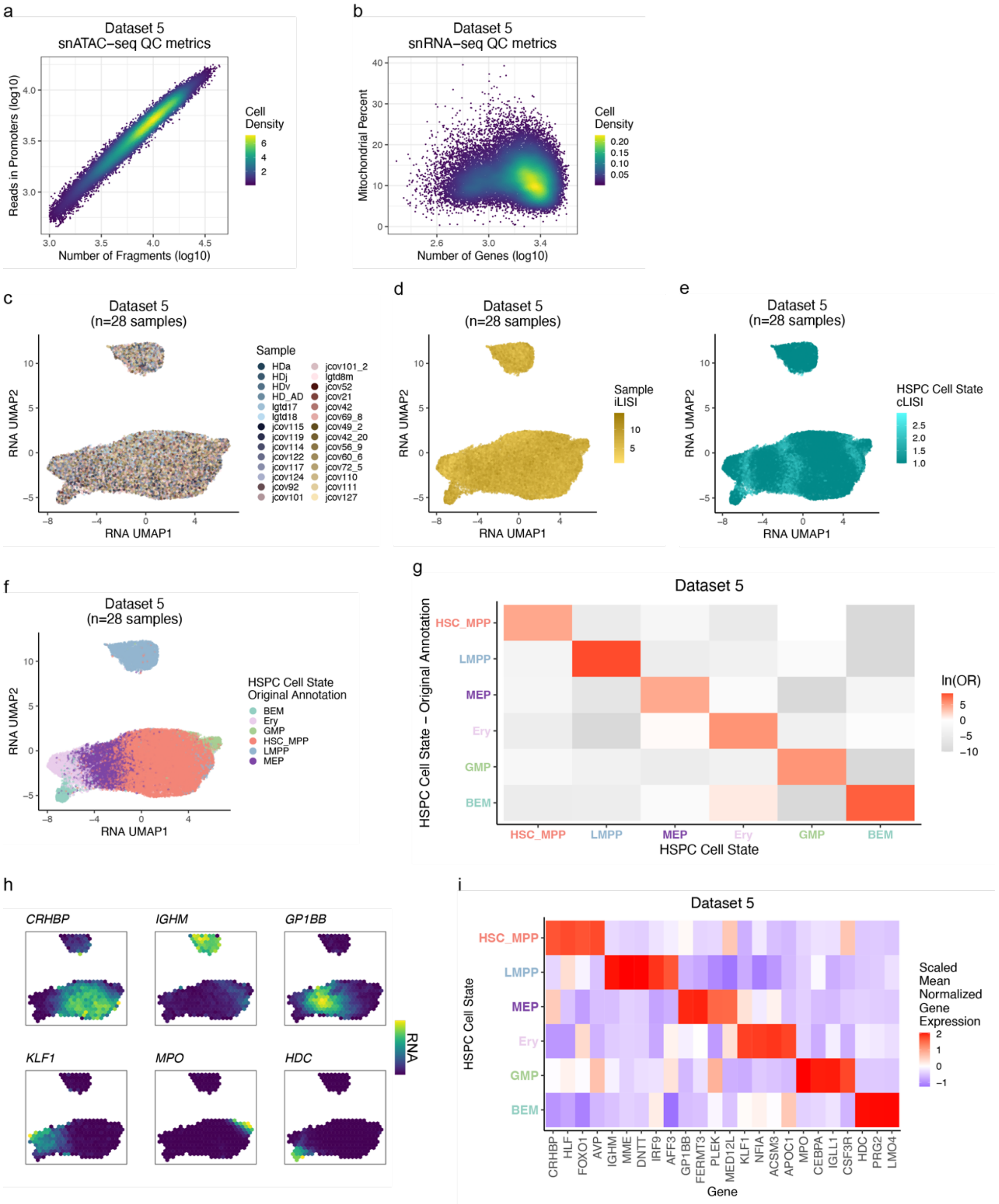

**Fig. S5.** Dataset 5. COVID-19 PBMC HSPC states.
**a.** snATAC-seq QC metrics of fragment count and reads in promoters, colored by cell density.
**b.** snRNA-seq QC metrics of gene count and mitochondrial percent, colored by cell density.
**c.** snRNA-seq UMAP colored by sample.
**d.** snRNA-seq UMAP colored by sample integration LISI (iLISI). Higher values (darker colors) are indictive of better batch correction.
**e.** snRNA-seq UMAP colored by cell type LISI (cLISI). Lower values (darker colors) are indictive of better bio-conservation.
**f.** snRNA-seq UMAP colored by HSPC states defined in the original study[22].
**g.** Natural log of the odds ratio between the re-annotated and original HSPC states from **f**. Non-

significant values (FDR>0.05) are white.

**h.** Binned mean-normalized marker gene expression on snRNA-seq UMAP; yellow denotes high expression.

**i**. Scaled mean-normalized marker gene expression across re-annotated HSPC states.

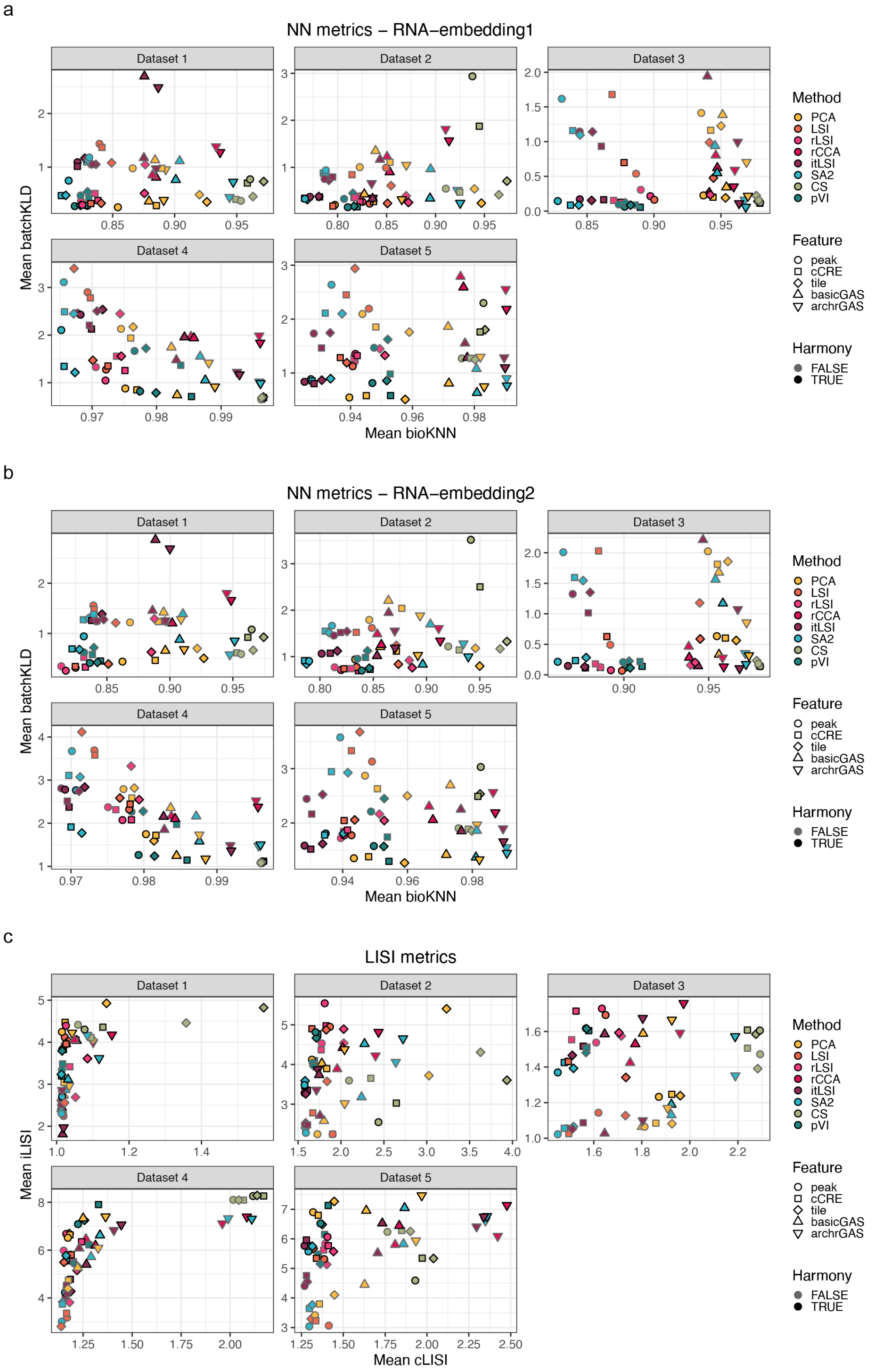


**Fig. S6.** Pipeline metric means across datasets.
Mean bio-conservation (**x-axis**) and batch correction (**y-axis**) metrics averaged across cells per dataset using **a.** NN metrics with RNA-embedding1, **b.** NN metrics with RNA-embedding2, and **c.** LISI metrics.

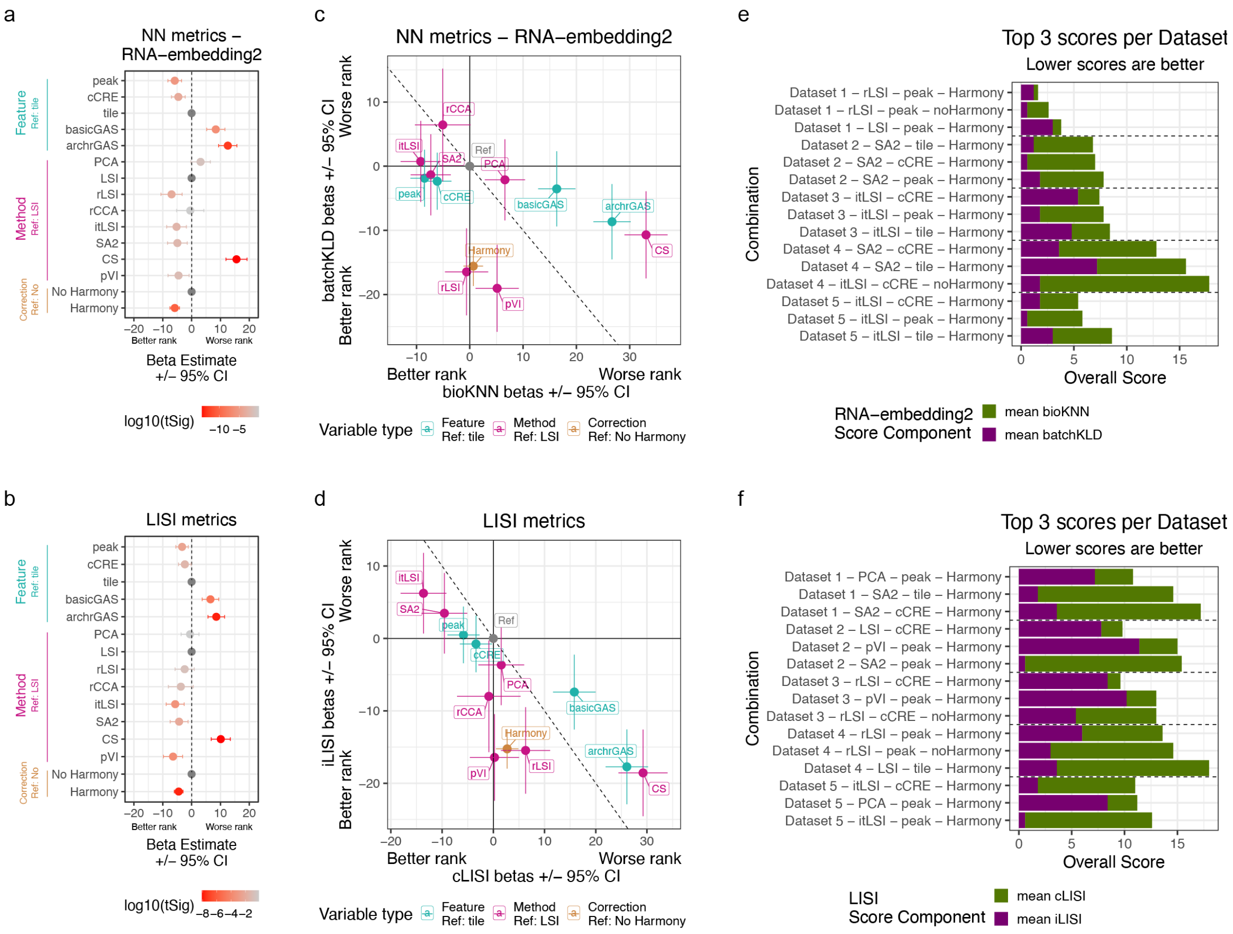


**Fig. S7.** Pipeline rankings with additional metrics.
**a.-b.** Combined linear model relating overall score (60% ranked mean bio-conservation metric + 40% ranked mean batch correction metric) to dataset, feature, method, and correction combinations for **a.** NN metrics using RNA-embedding2 and **b.** LISI metrics.
**c.-d.** Separated linear models as in **a.-b.**, but ranked mean bio-conservation (**x-axis**) vs ranked mean batch correction (**y-axis**) for **c.** NN metrics using RNA-embedding2 and **d.** LISI metrics.
**e.-f.** Top 3 overall scores per dataset, colored by ranked mean bio-conservation and ranked mean batch correction components.

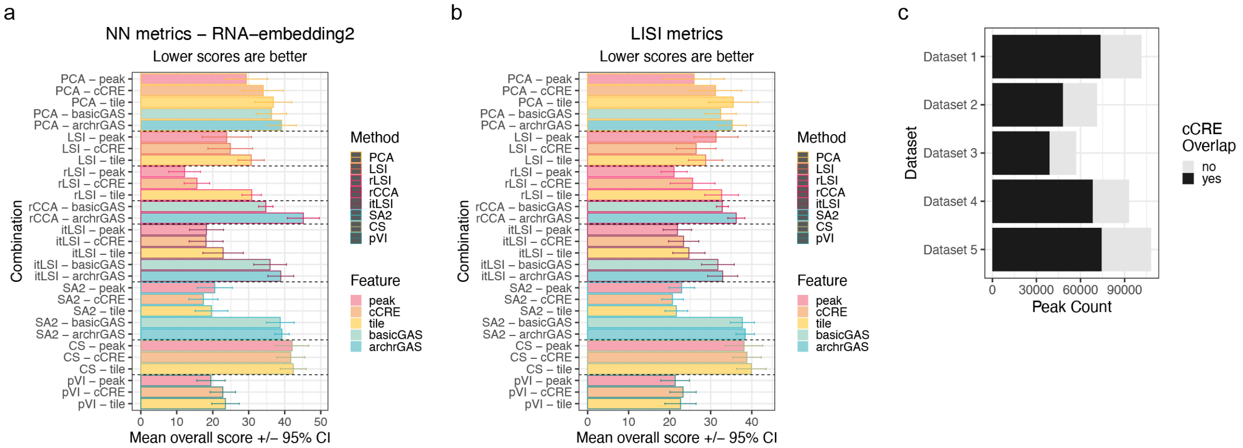


**Fig. S8.** ATAC features outperformed GAS features across metrics.
**a.-b.** The mean overall scores (60% bio-conservation rank + 40% batch correction rank) across datasets and correction choice for each method and feature combination for **a.** RNA-embedding2 NN metrics and **b.** LISI metrics. Lower scores are better. Error bars are the 95% confidence interval (CI).
**c.** The number of peaks per dataset overlapping or not overlapping a cCRE feature.

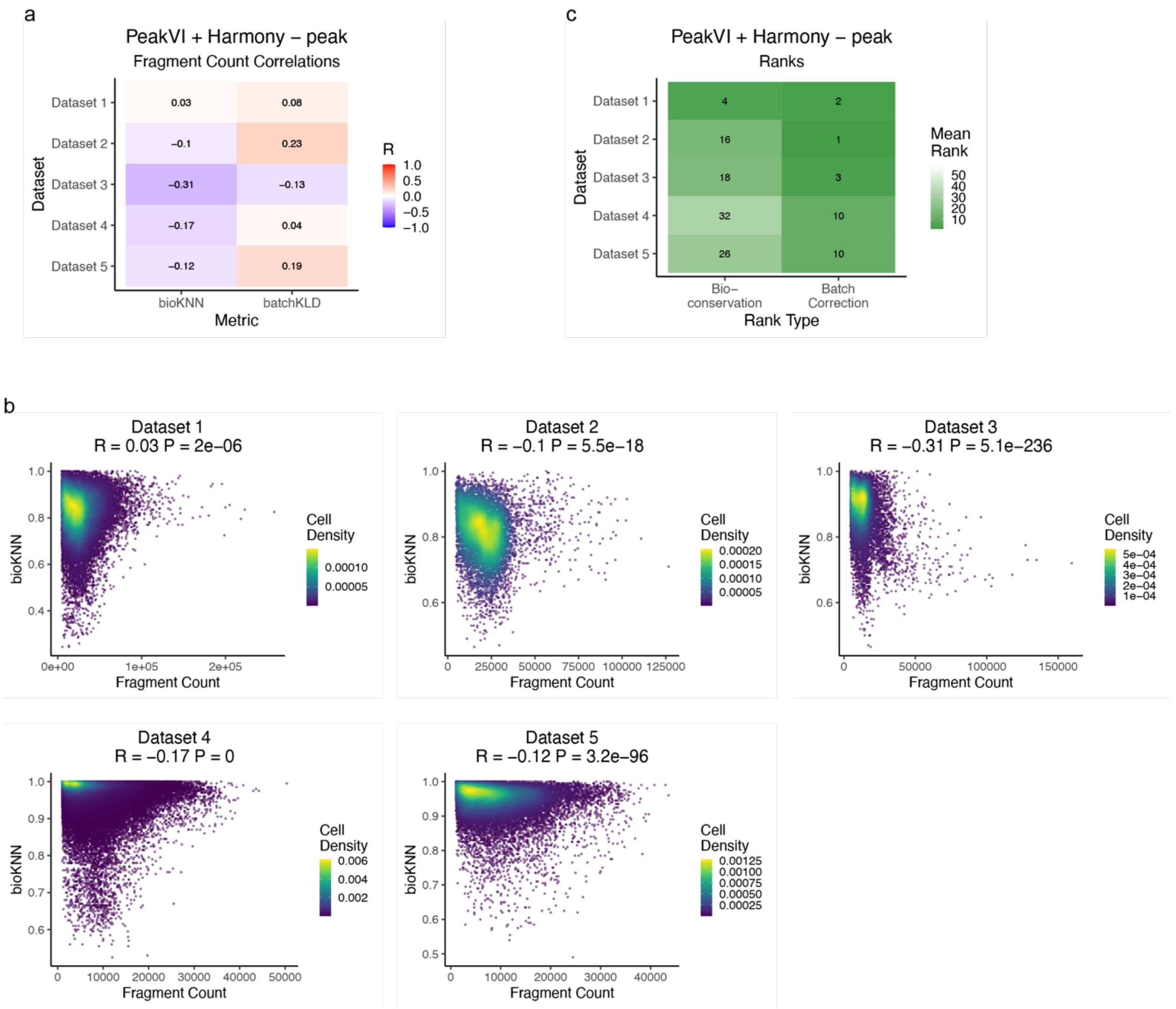


**Fig. S9.** PeakVI fragment bias affects bioKNN metrics and bio-conservation rankings.

**a.** Pearson correlations between fragment count and RNA-embedding1 NN metrics for PeakVI + Harmony with peak features across datasets.

**b.** Scatterplot between fragment count and bioKNN for PeakVI + Harmony with peak features across datasets. Cells are colored by cell density within each dataset. Pearson correlation estimates and p-values shown.

**c.** Bio-conservation and batch correction rankings for PeakVI + Harmony with peak features across datasets. Ranks span 1 (best) to 58 (worst).

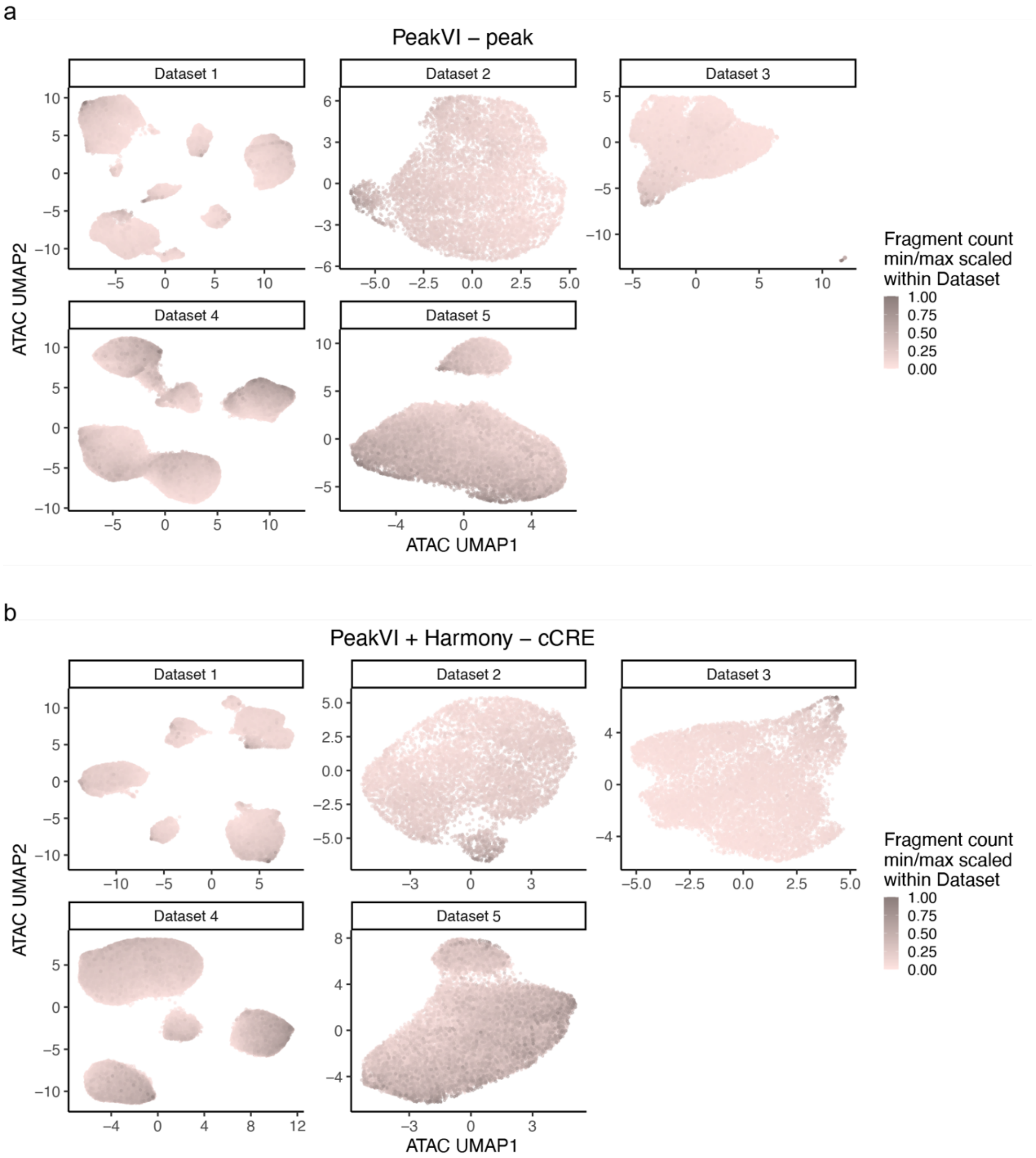


**Fig. S10.** PeakVI biased by fragment count across correction and features.
snATAC-seq UMAPs colored by fragment count for all Benchmarking Datasets using **a.** PeakVI with peak features and **b.** PeakVI + Harmony with cCRE features.

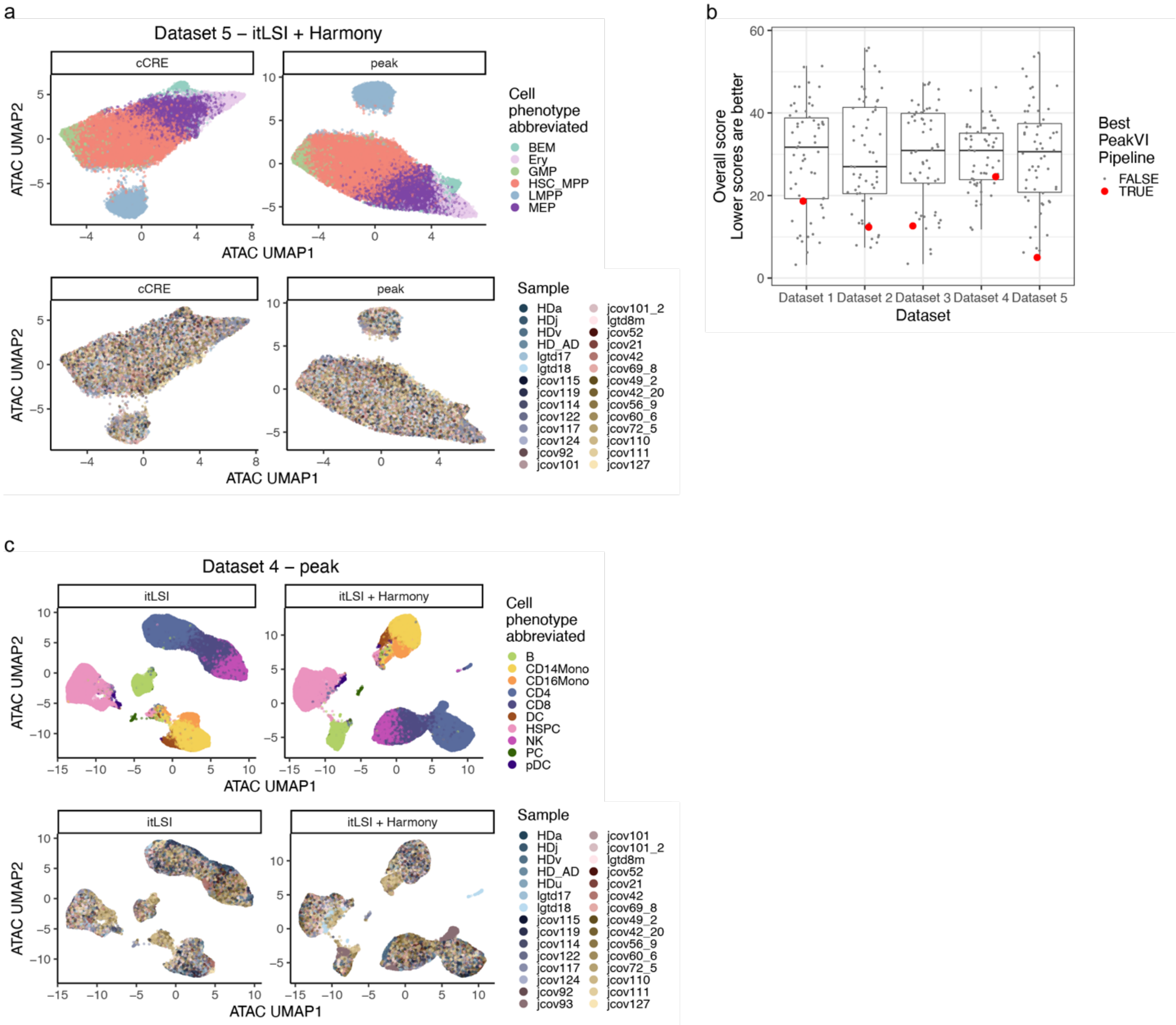


**Fig. S11.** ArchR itLSI preferred cell states over cell types.
**a.** snATAC-seq UMAPs colored by cell phenotype (**top**) and sample (**bottom**) for Dataset 5 using ArchR itLSI + Harmony with cCRE (**left**) or peak (**right**) features.
**b.** Overall scores of all 58 pipelines for each Benchmarking Dataset. The best PeakVI pipeline in each Dataset is highlighted: Dataset 1 – cCRE with Harmony, Dataset 2 – peak with Harmony, Dataset 3 – tile with Harmony, Dataset 4 – peak with Harmony, Dataset 5 – peak with Harmony.
**c.** snATAC-seq UMAPs colored by cell phenotype (**top**) and sample (**bottom**) for Dataset 4 with peak features using ArchR itLSI (**left**) or ArchR itLSI + Harmony (**right**).

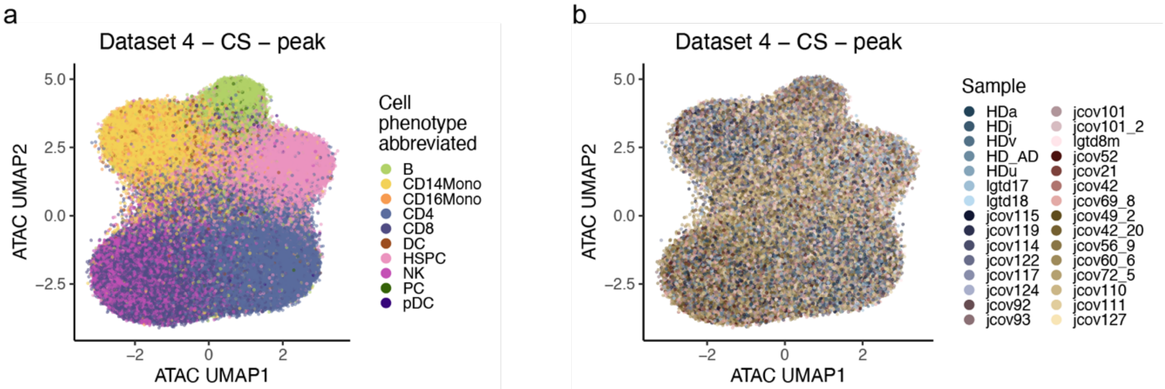


**Fig. S12.** CellSpace had poor bio-conservation yet good batch correction.
snATAC-seq UMAPs colored by **a.** cell phenotype and **b.** sample for Dataset 4 using CellSpace with peak features. This was the top performing pipeline that used CellSpace across Benchmarking Datasets.

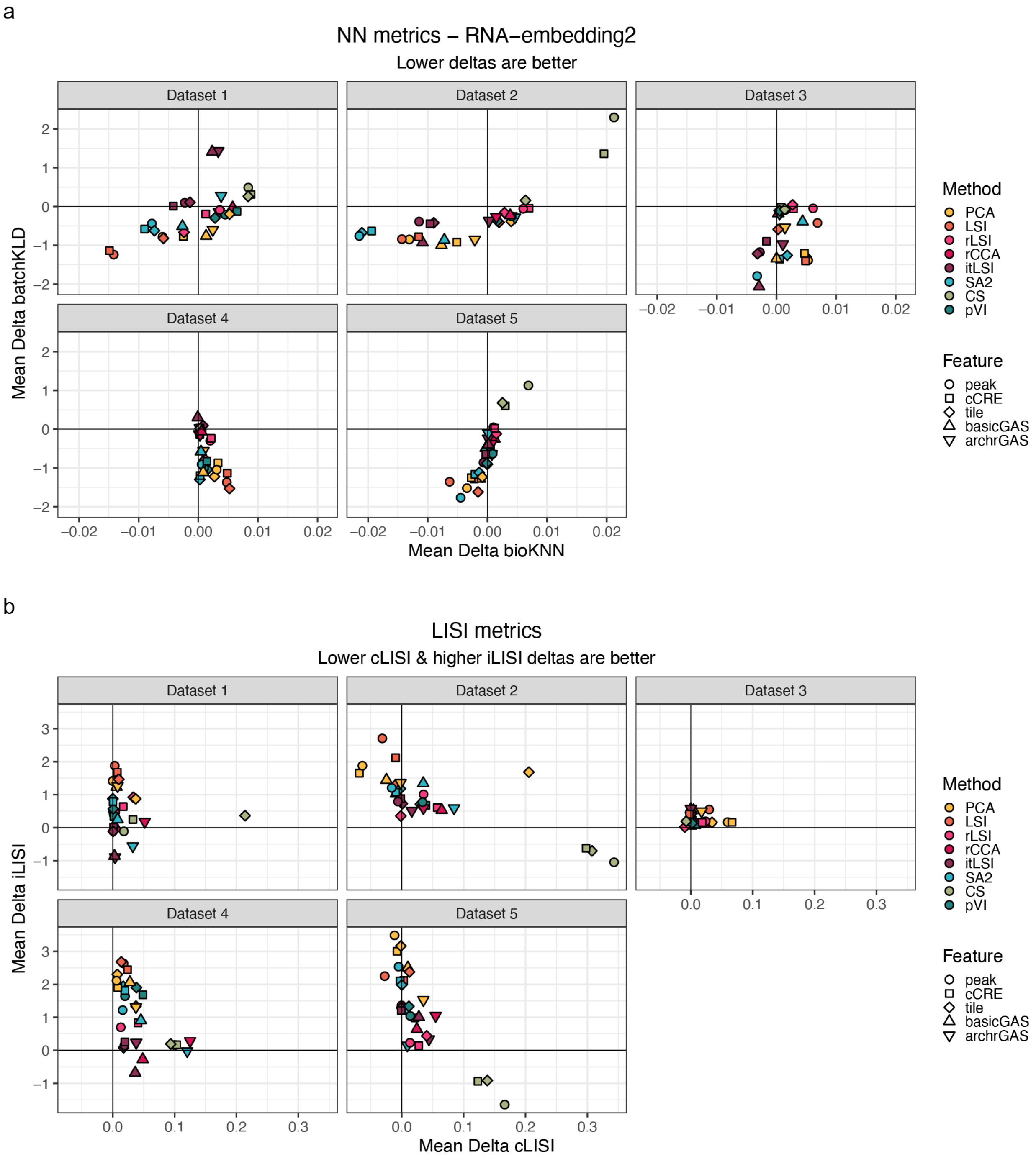


**Fig. S13.** Harmony Deltas across metrics.
Mean bio-conservation (**x-axis**) and batch correction (**y-axis**) Harmony deltas across method/feature combinations in all Benchmarking Datasets for **a.** RNA-embedding2 NN metrics and **b.** LISI metrics. Harmony deltas were calculated as the Harmony metrics subtracted by the no Harmony metrics for the same method/feature combination.

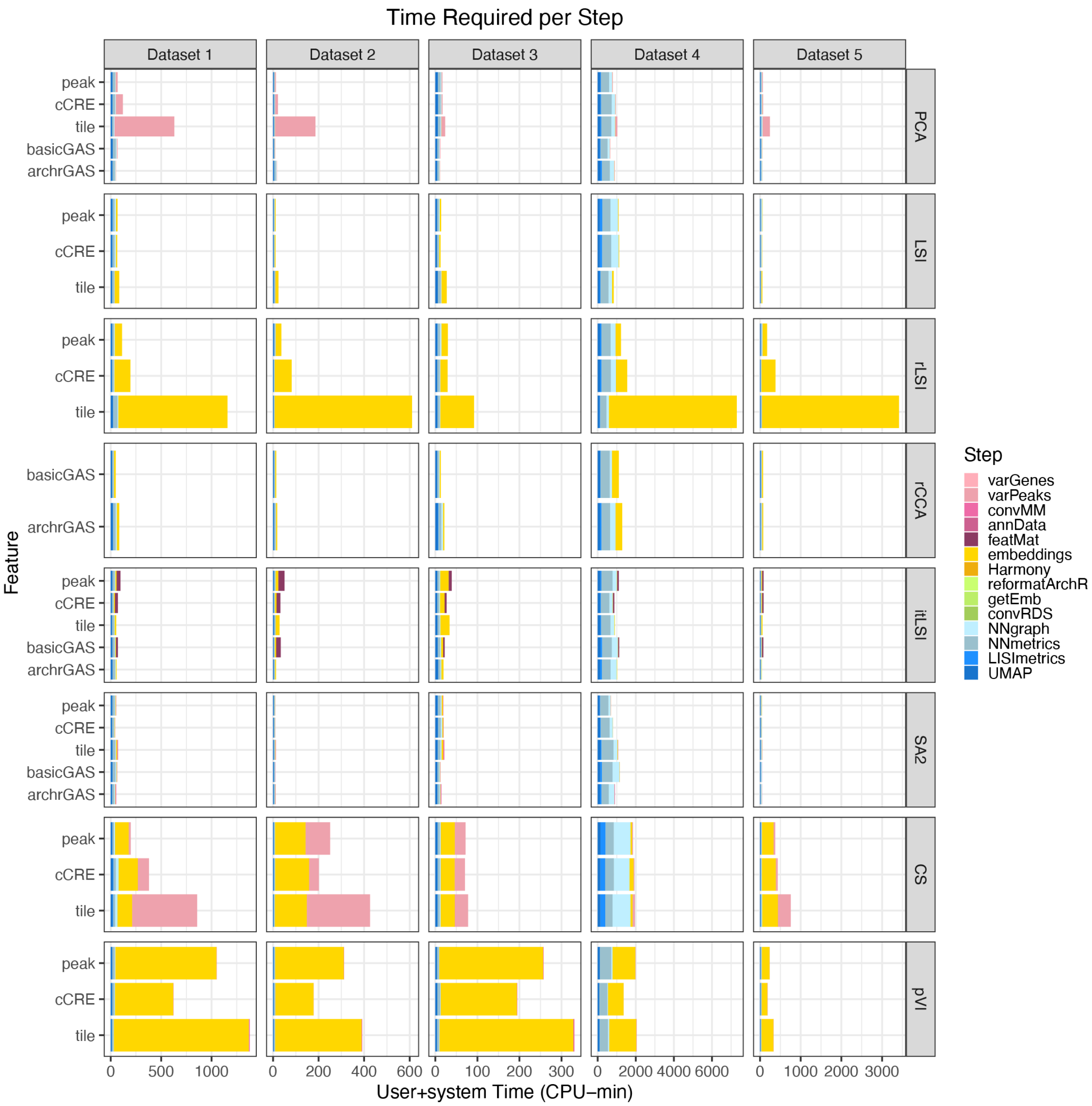


**Fig. S14.** Time requirement per pipeline step.
User+system time requirements per step across Benchmarking Datasets and Methods. Individual steps spanned pre-processing (pinks), embedding generation (yellow), Harmony correction (gold), post-processing (greens), and metric/visualization (blues). Values calculated with /usr/bin/time (**Methods**).

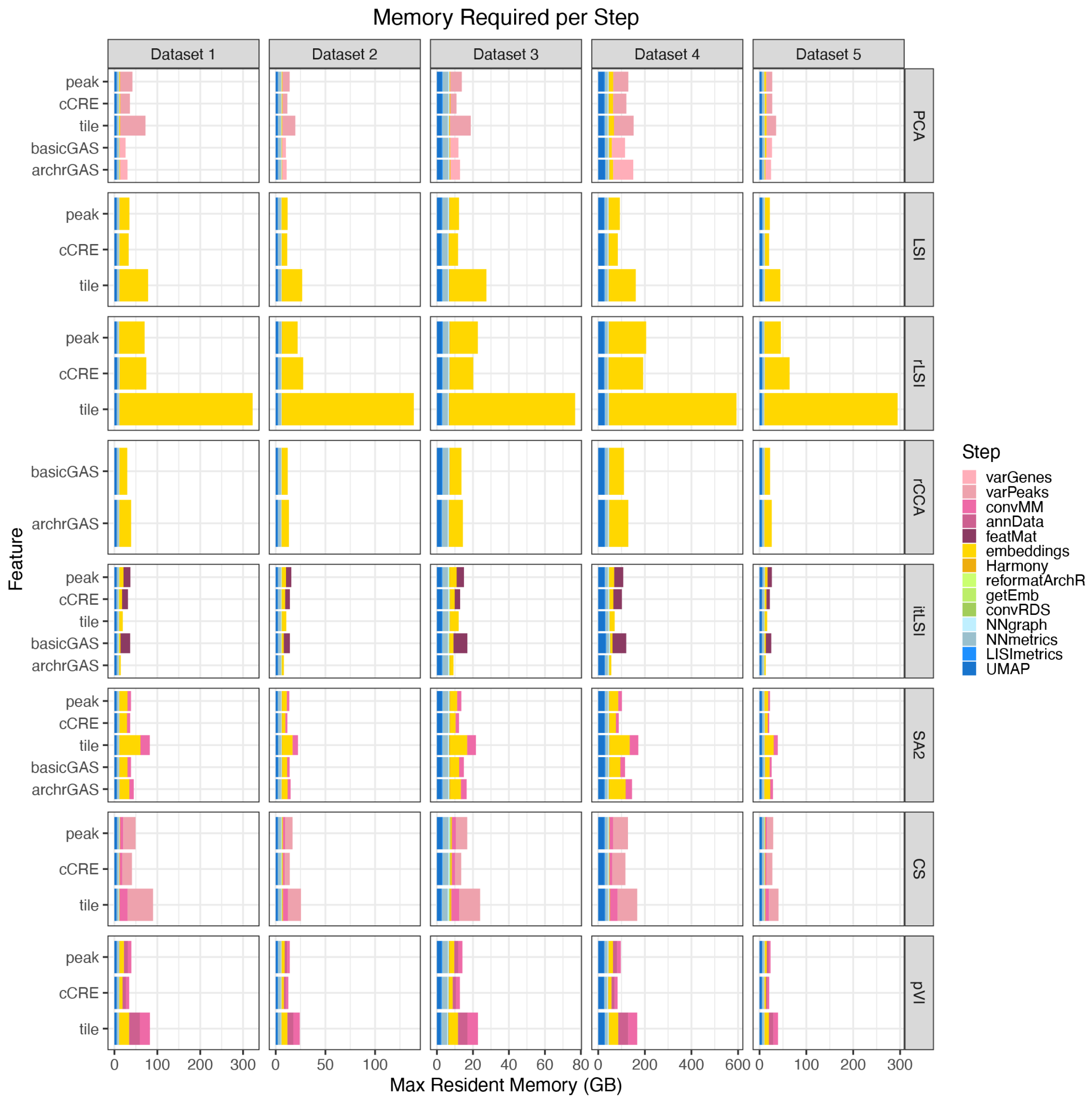


**Fig. S15.** Memory requirement per pipeline step.
Max resident memory requirements per step across Benchmarking Datasets and Methods. Individual steps spanned pre-processing (pinks), embedding generation (yellow), Harmony correction (gold), post-processing (greens), and metric/visualization (blues). Values calculated with /usr/bin/time (**Methods**).

a

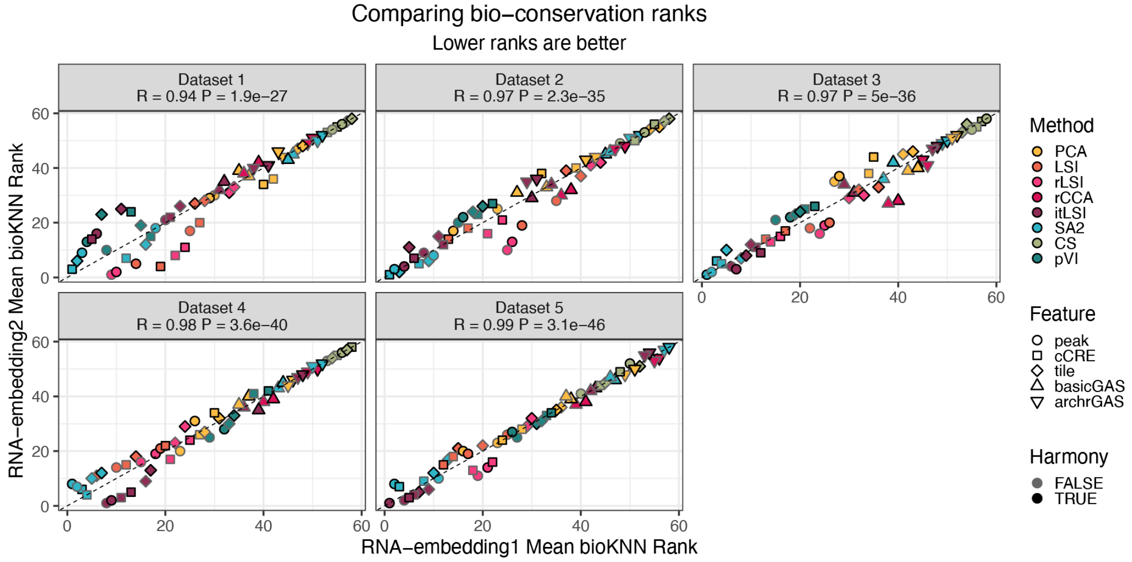


b

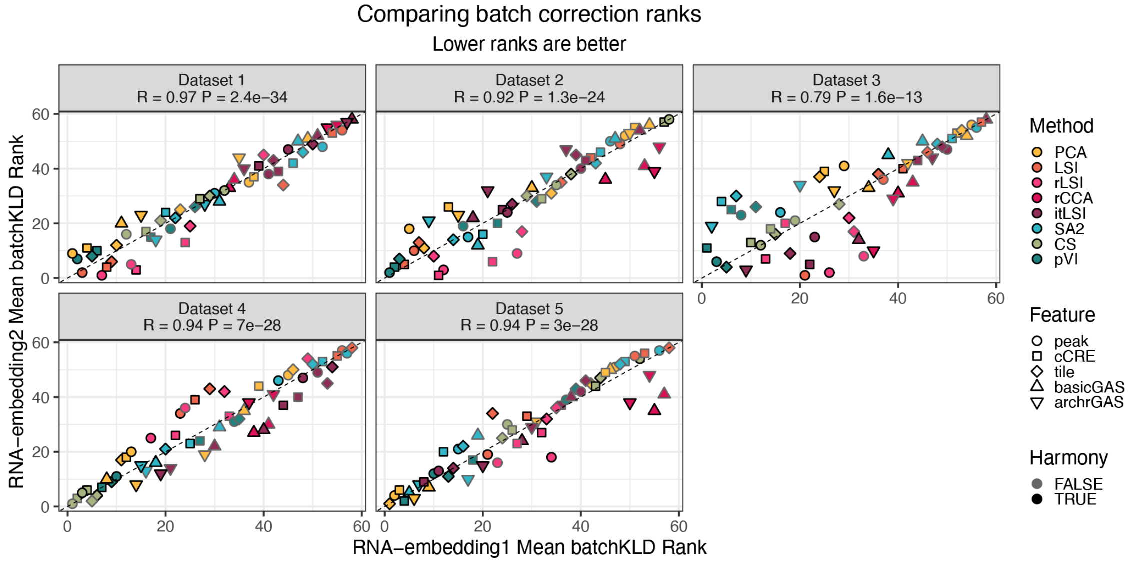


**Fig. S16.** RNA-embedding1 vs RNA-embedding2 NN metrics.
**a.** Bio-conservation rankings of mean bioKNN metrics using RNA-embedding1 (**x-axis**) vs RNA-embedding2 (**y-axis**) across datasets. Lower ranks were better. Pearson correlation estimates and p-values shown.
**b.** Batch correction rankings of mean batchKLD metrics using RNA-embedding1 (**x-axis**) vs RNA-embedding2 (**y-axis**) across datasets. Lower ranks were better. Pearson correlation estimates and p-values shown.

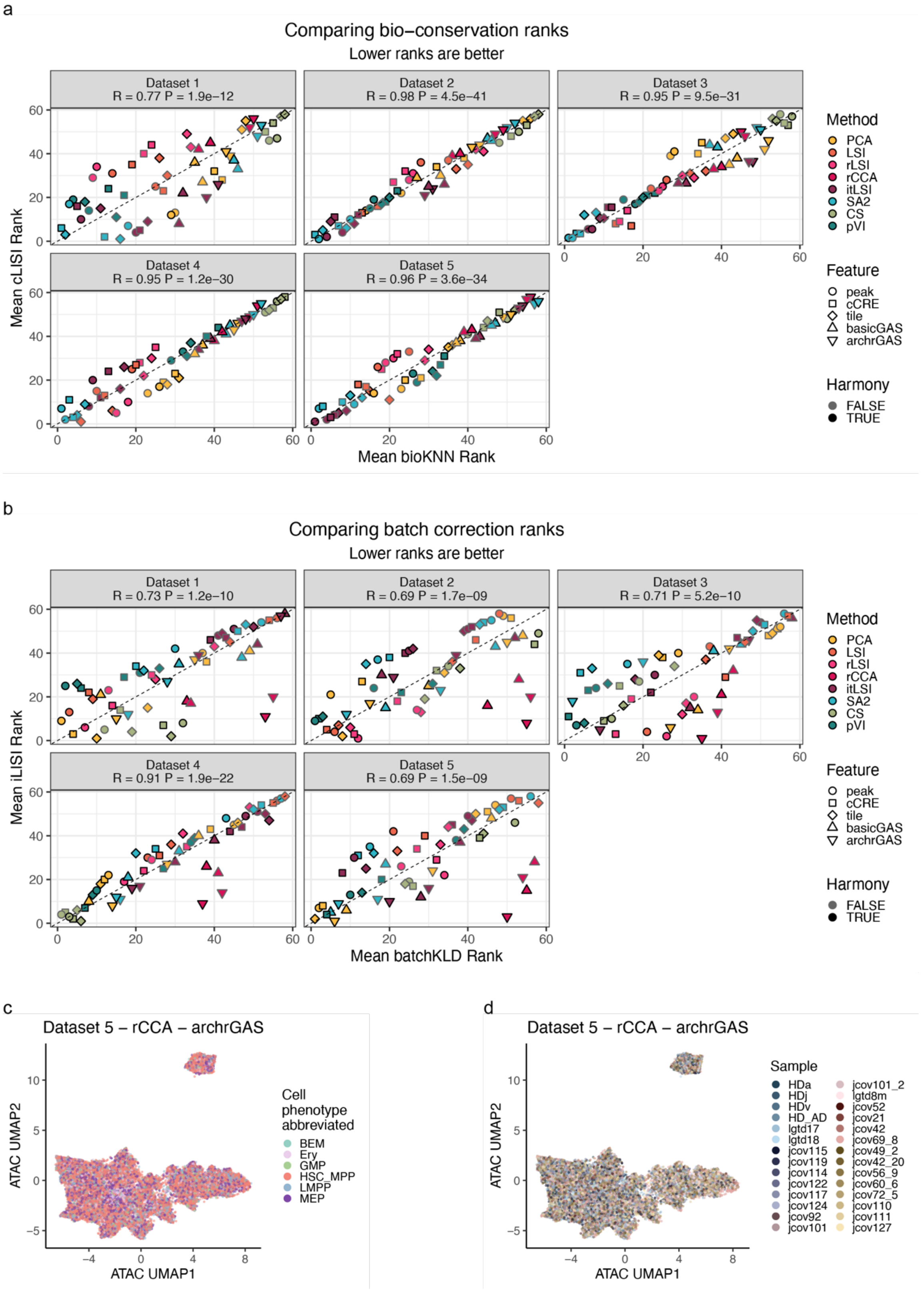


**Fig. S17.** RNA-embedding1 NN metrics vs LISI metrics.
**a.** Bio-conservation rankings of mean RNA-embedding1 bioKNN metrics (**x-axis**) by mean cLISI

metrics (**y-axis**) across datasets. Lower ranks were better. Pearson correlation estimates and p-values shown.
**b.** Batch correction rankings of mean RNA-embedding1 batchKLD metrics (**x-axis**) by mean iLISI metrics (**y-axis**) across datasets. Lower ranks were better. Pearson correlation estimates and p-values shown.
**c.-d.** snATAC-seq UMAPs colored by **c.** cell phenotype and **d.** sample for Dataset 5 using rCCA with archrGAS features.

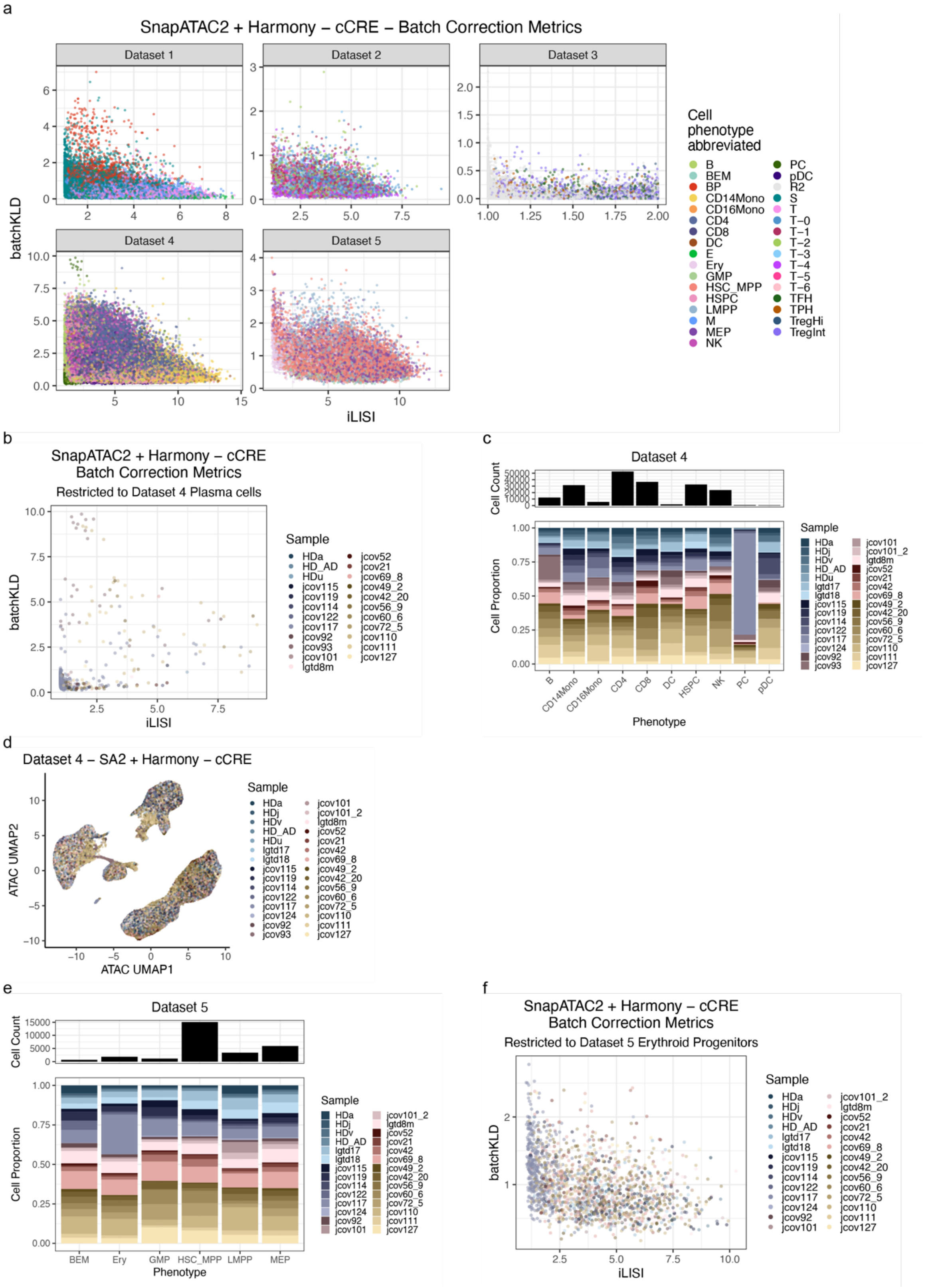


**Fig. S18.** SnapATAC2 exemplified batchKLD strengths over iLISI.
**a.** RNA-embedding1 batchKLD (**y-axis**) vs iLISI (**x-axis**) metrics from SnapATAC2 + Harmony

cCRE pipeline for all Benchmarking Datasets.
**b.** RNA-embedding1 batchKLD (**y-axis**) vs iLISI (**x-axis**) metrics from SnapATAC2 + Harmony cCRE pipeline subset to Dataset 4 plasma cells.
**c.** The total cell counts across all Dataset 4 cell phenotypes (**top**) as well as the proportion of cells per sample (**bottom**).
**d.** snATAC-seq UMAPs colored by sample for Dataset 4 cells using SnapATAC2 + Harmony with cCRE features.
**e.** The total cell counts across all Dataset 5 cell phenotypes (**top**) as well as the proportion of cells per sample (**bottom**).
**f.** RNA-embedding1 batchKLD (**y-axis**) vs iLISI (**x-axis**) metrics from SnapATAC2 + Harmony cCRE pipeline subset to Dataset 5 Erythroid progenitors.
